# Primary exposure to SARS-CoV-2 variants elicits convergent epitope specificities, immunoglobulin V gene usage and public B cell clones

**DOI:** 10.1101/2022.03.28.486152

**Authors:** Noemia S. Lima, Maryam Musayev, Timothy S. Johnston, Danielle A. Wagner, Amy R. Henry, Lingshu Wang, Eun Sung Yang, Yi Zhang, Kevina Birungi, Walker P. Black, Sijy O’Dell, Stephen D. Schmidt, Damee Moon, Cynthia G. Lorang, Bingchun Zhao, Man Chen, Kristin L. Boswell, Jesmine Roberts-Torres, Rachel L. Davis, Lowrey Peyton, Sandeep R. Narpala, Sarah O’Connell, Jennifer Wang, Alexander Schrager, Chloe Adrienna Talana, Kwanyee Leung, Wei Shi, Rawan Khashab, Asaf Biber, Tal Zilberman, Joshua Rhein, Sara Vetter, Afeefa Ahmed, Laura Novik, Alicia Widge, Ingelise Gordon, Mercy Guech, I-Ting Teng, Emily Phung, Tracy J. Ruckwardt, Amarendra Pegu, John Misasi, Nicole A. Doria-Rose, Martin Gaudinski, Richard A. Koup, Peter D. Kwong, Adrian B. McDermott, Sharon Amit, Timothy W. Schacker, Itzchak Levy, John R. Mascola, Nancy J. Sullivan, Chaim A. Schramm, Daniel C. Douek

## Abstract

An important consequence of infection with a SARS-CoV-2 variant is protective humoral immunity against other variants. The basis for such cross-protection at the molecular level is incompletely understood. Here we characterized the repertoire and epitope specificity of antibodies elicited by Beta, Gamma and ancestral variant infection and assessed their cross-reactivity to these and the more recent Delta and Omicron variants. We developed a high-throughput approach to obtain immunoglobulin sequences and produce monoclonal antibodies for functional assessment from single B cells. Infection with any variant elicited similar cross-binding antibody responses exhibiting a remarkably conserved hierarchy of epitope immunodominance. Furthermore, convergent V gene usage and similar public B cell clones were elicited regardless of infecting variant. These convergent responses despite antigenic variation may represent a general immunological principle that accounts for the continued efficacy of vaccines based on a single ancestral variant.

## Main Text

Over the course of the SARS-CoV-2 pandemic, selective immune pressure is proposed to have led to the accumulation of changes in residues targeted for antibody recognition and neutralization, most importantly in the receptor binding domain (RBD)^1, 2^. While CD4 and CD8 T cell responses do not seem to be substantially impacted by variant substitutions^3^, neutralizing capacity and some Fc-mediated functionality of antibodies induced by the ancestral SARS-CoV-2 variant (WA1) are significantly reduced against later variants^4, 5^. Despite this, first generation vaccines based on the WA1 sequence continue to provide protection from severe disease and death^6^ even against antigenically distant variants such as Delta (PANGO lineage B.1.617.2) and Omicron (B.1.1.529). The mechanism of this cross-protection is not fully understood at the molecular level, even though the humoral response to the ancestral virus has been well characterized^7, 8, 9, 10^. Notably, the response to ancestral WA1 is highly consistent and includes polarization toward specific IG V_H_ genes^11, 12, 13^ and convergent V(D)J rearrangements (“public clones”) found in multiple individuals^13, 14, 15^. High-resolution analysis of the immune responses to other, antigenically divergent, variants may be leveraged to explore the extent of conservation of these responses and to shed light on mechanisms of cross-protection. In a cohort of convalescent individuals infected with WA1, Beta (B.1.351), or Gamma (P.1), we use a novel method for high-throughput, cloning-free recombinant mAb synthesis and sequencing to investigate epitope targeting, V_H_ gene usage, and B cell clonal repertoires against these variants as well as Delta and Omicron.

We collected serum or plasma and PBMC from individuals infected with WA1, Beta, or Gamma variants at 17-38 days after symptom onset (Extended Data Fig. 1) to compare antibody and B cell responses. All individuals were previously naïve to SARS-CoV-2. To focus on the total antigen-specific B cell repertoire, we selected samples from early convalescence, when frequencies of B and T cells are typically high, irrespective of neutralization titers.

We measured serum binding titers to variant spike (S) protein expressed on the surface of HEK293T cells (Fig. 1A) and to soluble stabilized variant S trimers (S-2P) and RBD using a Meso Scale Discovery electrochemiluminescence immunoassay (MSD-ECLIA) (Extended Data Fig. 2A). Both assays showed that all convalescent individuals had antibodies against the homologous S as well as cross-reactive antibodies to S from other variants. The WA1-infected individuals showed a significant reduction in antibody titers binding to Omicron BA.1 S (Fig. 1A) and to Beta RBD (Extended Data Fig. 2A). The Beta-infected individuals exhibited the highest titers against Beta S and significantly reduced titers against D614G, Delta and Omicron BA.1 (Fig. 1A). The Gamma-infected individuals showed the least variation in antibody binding titers across the different variants (Fig 1A). Consistent with previous reports^16, 17^, variant-infected individuals recognized WA1 RBD at similar levels as the homologous RBD (Extended Data Fig. 1A). Individuals with the highest serum binding titers (SAV1, SAV3 and A49) could cross-neutralize WA1, Beta and Gamma, and showed lower potency against Delta and Omicron BA.1 and BA.2 variants (Fig. 1B). Other individuals completely lost neutralization against Delta and Omicron variants, except for SAV11 who retained a low neutralization titer against Omicron BA.2 (Fig. 1B).

**Fig. 1:**
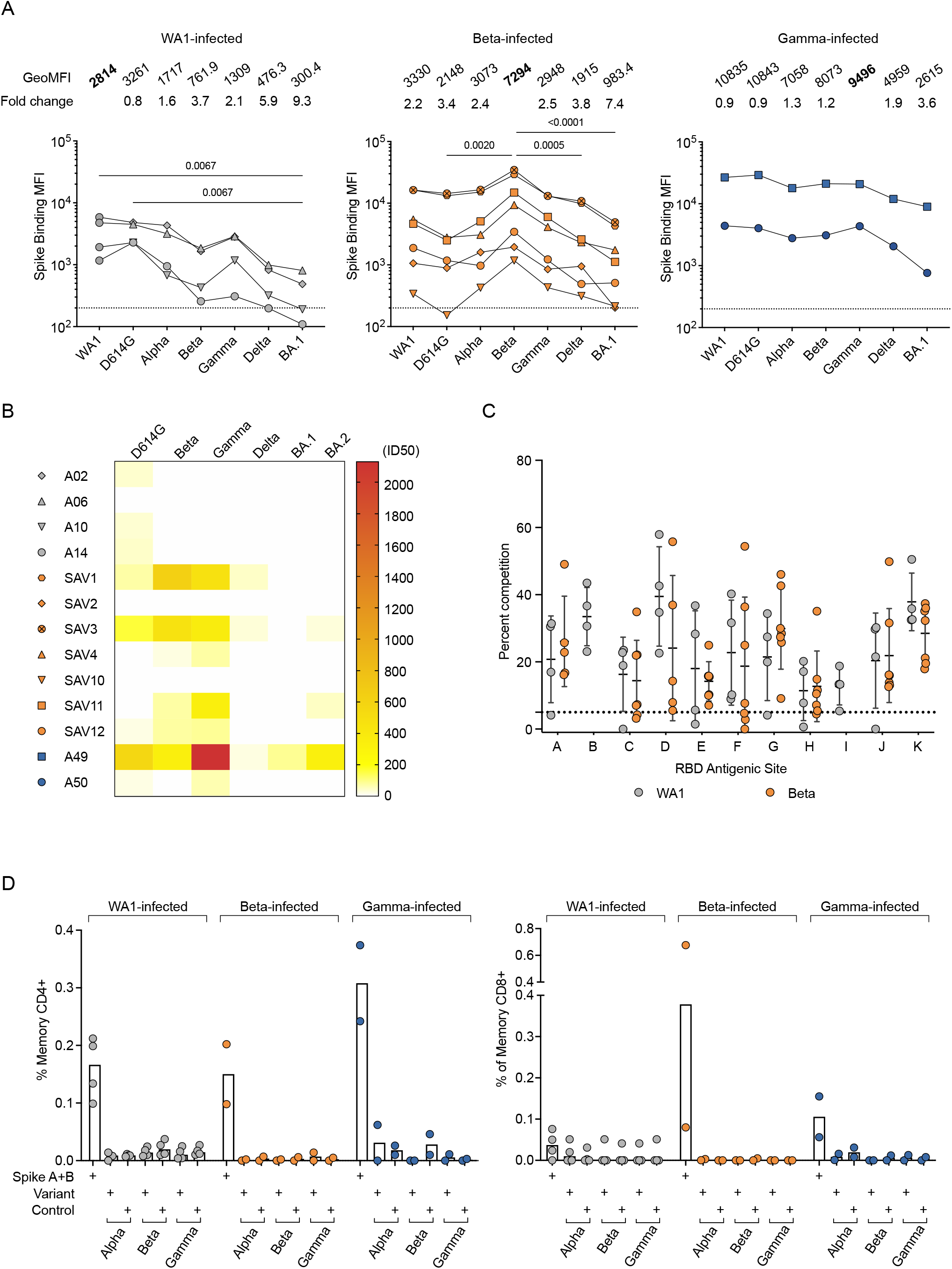
Homologous and cross-reactive antibodies induced by WA1 and variant infections. (**A**) Antibody binding titers against multiple variants assessed by cell surface binding assay. (**B**) Heatmap showing neutralizing antibody titers (reciprocal 50% inhibitory dilution) for each individual labeled on the left against each variant indicated on the top. (**C**) Epitope mapping on homologous spike by competition assay using surface plasmon resonance. Antibodies CB6 (RBD-B epitope) and A19-30.1 (RBD-I) do not bind to Beta and competition is not measured at these sites. (**D**) CD4 (left) and CD8 (right) T cell responses to WA1 spike peptide pools A+B, selected pools containing altered variant peptides and control pool containing correspondent peptides for each variant pool.

We next used a surface plasmon resonance (SPR)-based competition assay^18, 19^ to characterize epitopes targeted by serum antibodies (Extended Data Fig. 2B). Notably, when the binding activity of each serum was characterized against the homologous S, the patterns of reactivity were comparable between individuals infected either with WA1 or Beta (Fig. 1C), revealing a conserved immunodominance hierarchy across variants, despite antigenic changes. Likewise, there were no differences in competition at each epitope when sera from Beta- or Gamma-infected individuals were mapped against WA1, Beta, or Delta S (Extended Data Fig. 2, C and D).

We evaluated the ability of T cells elicited by Beta and Gamma infections to recognize WA1 S peptides by measuring upregulation of CD69 and CD154 on CD4 T cells, and production of IFN-γ, TNF, or IL-2 by CD8 T cells (Extended Data Fig. 2E). CD4 and CD8 T cell responses to WA1 S peptides were similar in Beta- and Gamma-infected individuals compared to WA1-infected individuals (Fig. 1D). When stimulated with selected peptides covering only regions containing substitutions in each variant, CD4 and CD8 T cell responses were minimal, suggesting that the substituted residues are not included within immunodominant T cell epitopes (Fig. 1D).

The three individuals in our cohort with the highest binding titers (Fig. 1A) were selected for in-depth characterization of the antibody repertoire and identification of mAb binding patterns. We developed a method for rapid assembly, transfection, and production of immunoglobulins (abbreviated to RATP-Ig) from single-sorted B cells. RATP-Ig relies on 5’-RACE and high-fidelity DNA assembly to produce recombinant heavy and light chain-expressing linear DNA cassettes, which can be directly transfected into 96-well microtiter mammalian cell cultures. Resulting culture supernatants containing the expressed mAbs can then be tested for functionality (Extended Data Fig. 3). We sorted cross-reactive WA1^+^Beta^+^ B cells (Extended Data Fig. 4, A-C) from the three selected individuals, resulting in a total of 509 single cells for analysis (Fig. 2A). We recovered paired heavy and light chain sequences from 355 (70%) of cells (Fig. 2A). In parallel, we screened the RATP-Ig supernatants by ELISA for binding to S-2P, RBD, and NTD derived from each of WA1, Beta, Gamma, and Delta variants, as well as S-2P from the Omicron variant (B.1.1.529). IgG binding at least one antigen was produced in 255 wells (50%) containing a B cell (Fig. 2, A and B). All three individuals yielded high levels of cross-reactive antibodies to S, NTD, and RBD (Fig. 2B and Supplementary Tables 1-3). Antibodies isolated from Beta-infected individuals SAV1 and SAV3 showed similar binding profiles, being dominated by cross-reactive mAbs among WA1, Beta, Gamma, and Delta variants. About half of these antibody populations comprised S-2P-only binding antibodies, with lower proportions binding NTD or RBD epitopes (Fig. 2B). From Gamma-infected individual A49, we recovered a population of mAbs that was dominated by RBD binders. While most antibodies isolated from individual A49 were also cross-reactive, we isolated a large number of mAbs whose epitope specificity we deemed indeterminate, appearing to bind both RBD and NTD (Fig. 2B and Supplementary Table 3), perhaps due to high background ELISA signal.

**Fig. 2:**
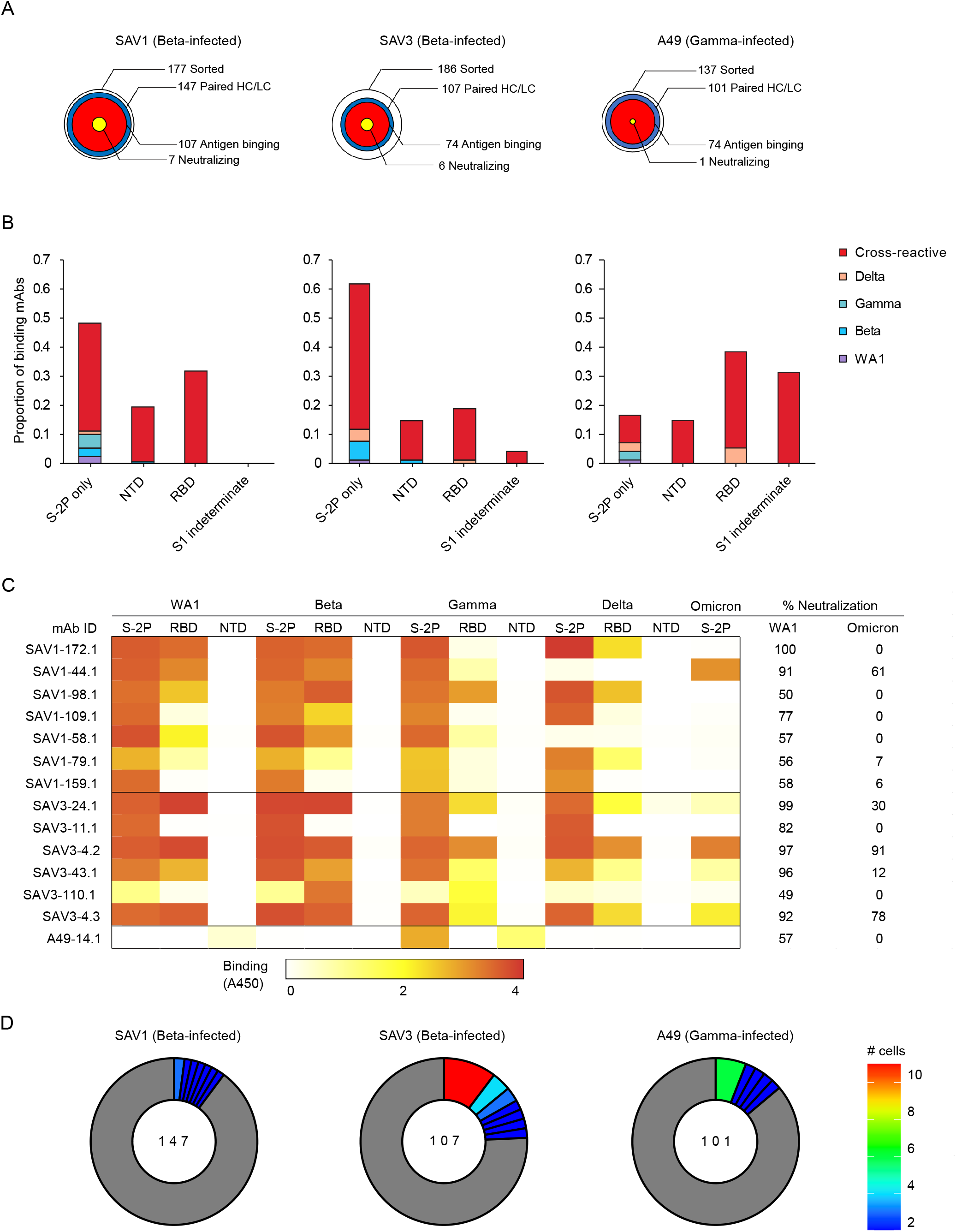
Functional Characterization of RATP-Ig Isolated mAbs. (**A**) RATP-Ig screening overviews for three individuals, represented as bullseyes. The area of each circle is proportional to the number of antibodies. (**B**) Supernatants were screened for antigen-specific binding by single-point ELISA for WA1, Beta, Gamma, and Delta S-2P, RBD, and NTD, as well as Omicron S-2P. Each panel represents data from a single individual, as in (A). (**C**) Neutralization screening of isolated antibodies at 4- or 6-fold supernatant dilutions using a D614G pseudovirus luciferase reporter assay, reported as % virus neutralized derived from reduction in luminescence. Associated ELISA heatmap reported as absorbance at 450nm (not quantitative). (**D**) Clonal expansion in each individual. Expanded clones are colored by the number of cells in each clone as shown on the right; singleton clones are shown in gray.

We next performed WA1 and Omicron pseudovirus neutralization screening for all supernatants at a 4- or 6-fold dilution. This assay identified 7, 6, and 1 antibodies neutralizing WA1 from individuals SAV1, SAV3, and A49, respectively (Fig. 2C). For most antibodies, neutralization ability was diminished when tested against Omicron pseudovirus. Only three antibodies (SAV1-44.1, SAV3-4.2 and SAV3-4.3) maintained greater than 50% Omicron pseudovirus neutralization at 4- or 6-fold dilution (Fig. 2C). Neutralizing antibodies were predominately cross-reactive and RBD-specific, except for two (SAV1-159.1 and SAV3-11.1) which bound to S-2P only and a single (A49-14.1) NTD-specific antibody (Fig. 2C). RBD-specific neutralizing antibodies were also the most potent of those isolated, with 6/12 neutralizing >90% of pseudovirus at 4-fold dilution. We validated the RATP-Ig results by selecting seven antibodies for heavy and light chain synthesis and expression and found RATP-Ig screening to be reliably predictive of mAb functionality, with 80/91 (88%) of functional interactions being reproduced (Extended Data Fig. 5). In summary, we found that primary infection with Beta or Gamma variants elicited similar cross-reactive B-cell responses, at single-cell resolution, targeting diverse SARS-CoV-2 epitopes.

While all three individuals had polyclonal antigen-specific repertoires (Fig. 2D), SAV3 and A49 had highly expanded clones matching a widely reported public clone using IGHV1-69 and IGKV3-11^9, 20, 21, 22, 23^. Members of this public clone were also recovered from SAV1, although they were not greatly expanded. RATP-Ig ELISA data indicated that these antibodies bound a non-RBD, non-NTD epitope on S-2P, consistent with available data for previously described members of this public clone. Notably, all but one of the antibodies we recovered from this public clone bound to Delta S-2P, and 11/17 also bound to Omicron S-2P. In addition, most antibodies from this public clone have been reported to bind SARS-CoV-1^9, 20, 21, 22, 23^, and one, mAb-123^21^, weakly binds endemic human coronaviruses HKU1 and 229E. We also found 2 antibodies, SAV1-109.1 and SAV1-168.1, with a YYDRxG motif in CDR H3 that can target the epitope of mAb CR3022 on RBD and produce broad and potent neutralization of a variety of sarbecoviruses^24^. While SAV1-168.1 was cross-reactive but non-neutralizing (Supplementary Table 1), SAV1-109.1 showed good neutralization potency and bound to WA1, Beta, Gamma and Delta, but not Omicron (Fig. 2C). Overall, 185 (90%) of the 206 WA-1/Beta cross-binding mAbs also bound Delta, while only 109 (53%) of those mAbs also bound Omicron (Supplementary Tables 1-3).

To investigate possible differences in targeting of domains outside of RBD, we further examined epitope specificities by flow cytometry (Extended Data Fig. 4, B and D). As expected, the frequency of antigen-specific cells generally correlated with serum binding titers, and cells capable of binding to heterologous variants were typically less frequent than those binding the infecting variant (Fig. 3A). In addition, both Beta- and Gamma-infected individuals showed higher frequencies of NTD-binding B cells against the homologous virus when compared to WA1-infected individuals (Fig. 3B).

**Fig. 3:**
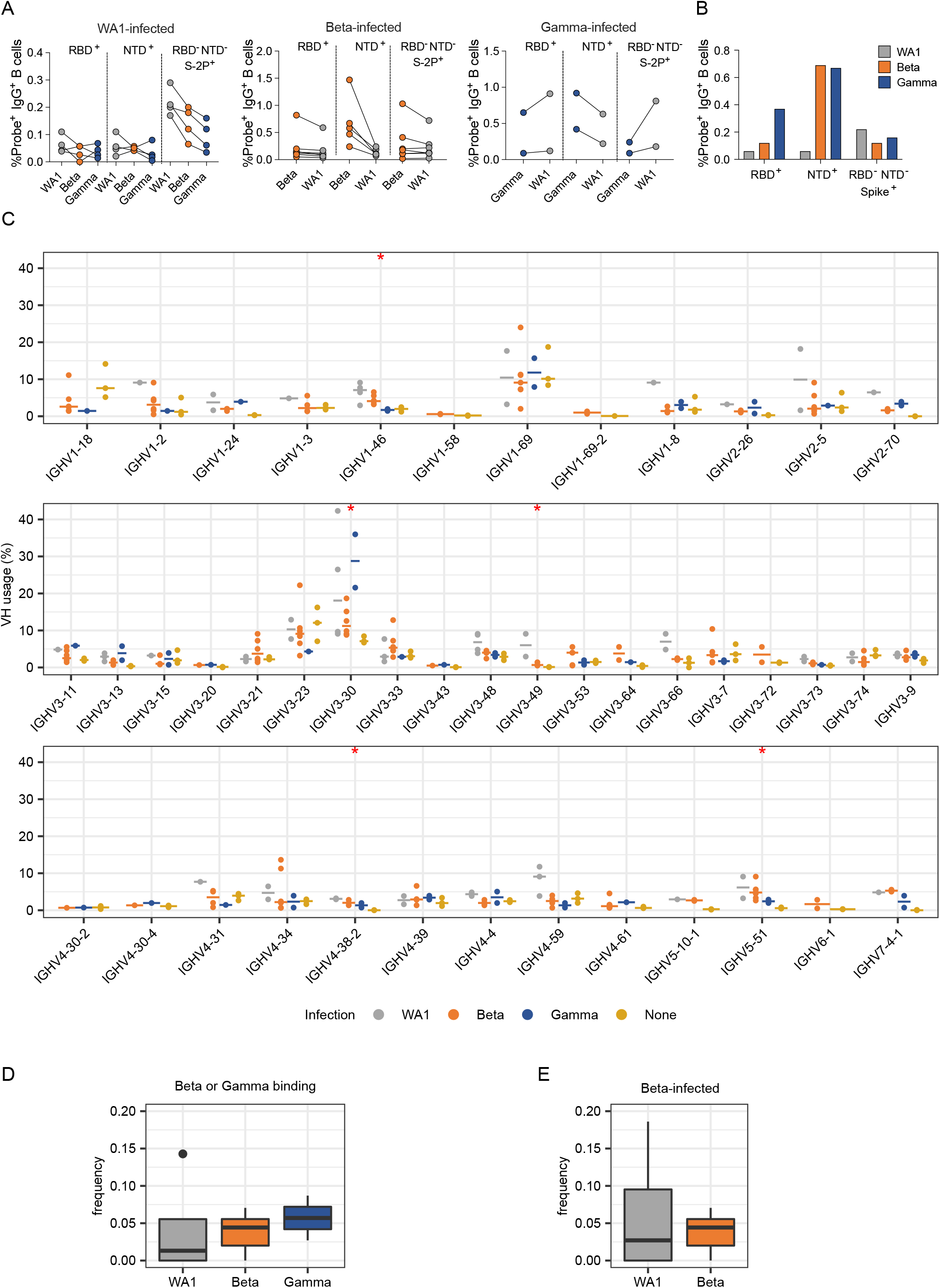
Anti-SARS-CoV-2 Ig repertoires. **(A)** Frequencies of probe^+^ B cells sorted for IG repertoire analysis. **(B)** Proportion of probe^+^ B cells binding to each domain. **(C)** SARS-CoV-2-specific VH repertoire analysis by infecting variant WA1, Beta and Gamma shown in grey, orange and blue, respectively, with data from pre-pandemic controls in yellow. X-axis shows all germline genes used; y-axis represents percent of individual gene usage. Stars indicate genes with at least one significant difference between groups; pairwise comparisons are in Extended Data Fig. 8. (**D**) and (**E**) Combined frequency of VH genes capable of giving rise to stereotypical Y501-dependent antibodies (IGHV4-30, IGHV4-31, IGHV4-39, and IGHV4-61) in (**D**) Beta- or Gamma-binding B cells from individuals infected with each variant or (**E**) B cells from Beta-infected individuals sorted with either WA1- or Beta-derived probes.

We generated libraries from sorted antigen-specific single cells using the 10x Genomics Chromium platform and recovered a total of 162, 319, and 107 paired heavy and light chain sequences from WA1-, Beta-, and Gamma-infected groups, respectively (Extended Data Figs. 4E and 6). As observed in the sequences identified via RATP-Ig, all three SARS-CoV-2-specific IG repertoires showed little clonal expansion. We then combined these data with the sequences generated by RATP-Ig for downstream analysis. Antigen-specific V gene usage was highly similar across all three infection types (Fig. 3C and Extended Data Fig. 7), with differences noted only for IGHV1-46 and IGLV1-47 (Extended Data Fig. 8). However, when we compared these antigen-specific repertories to the total memory B cell repertoire in pre-pandemic controls^25^, we observed significant enrichment for several genes (Fig. 3C and Extended Data Figs. 7 and 8). This highlights the convergence in responses to all SARS-CoV-2 variants we investigated.

Recent studies have shown that Y501-dependent mAbs derived from IGHV4-39 and related genes are overrepresented among neutralizing antibodies isolated from Beta-infected individuals^26, 27^. We therefore analyzed the observed frequency of these germline genes among Beta- and Gamma-binding B cells but found no significant differences based on infecting variant (Fig. 3D). Furthermore, we compared the frequency of sequences using these germline genes for WA1-versus Beta-binding B cells among Beta-infected individuals (excluding cross-reactive B cells isolated by RATP-Ig), and again found no difference in usage (Fig 3E). The lack of observed enrichment for these genes is likely due to the fact that neutralizing antibodies comprise only a small fraction of the antigen-specific binding repertoire^9, 28^, with the latter remaining highly conserved across individuals infected with different variants.

We next investigated SHM levels in these repertoires. The median V_H_ SHM levels among individuals was 0.3-6.6% in V_H_ and 0.0-3.0% in V_L_, compared to 6.7% and 2.4%, respectively, in the control repertoires. We then further examined SHM by both infecting variant and the probes used to isolate each cell. We found no differences in SHM in single probe-binding repertoires for either WA1- or Gamma-infected individuals (Fig. 4). Surprisingly, cross-reactive (WA1 and Beta) cells sorted for RATP-Ig had lower SHM than the single probe-binding repertoires sorted for 10x Genomics and sequencing. This may suggest that Beta S-2P is a better immunogen, capable of stimulating naïve B cells that require less SHM to gain cross-reactivity. Moreover, single probe-binding Beta-specific B cells from Beta-infected individuals had significantly higher SHM (median of 4.9% in V_H_ and 2.7% in V_L_) compared to single probe WA1-binding cells from the same individuals (2.1% and 0.8%, respectively) (Fig. 4). Other studies have also suggested the possibility that the immune response to Beta may be somewhat distinct from that against other SARS-CoV-2 variants, with neutralization appearing to wane more slowly and rising to higher levels after additional vaccine doses^18, 19^. Overall, the low levels of SHM across all the SARS-CoV-2-specific B cells that we isolated is consistent with prior reports^13, 22, 28^ and further demonstrates that the human immune system can readily generate antibodies capable of cross-binding multiple variants, regardless of infecting variant.

**Fig. 4:**
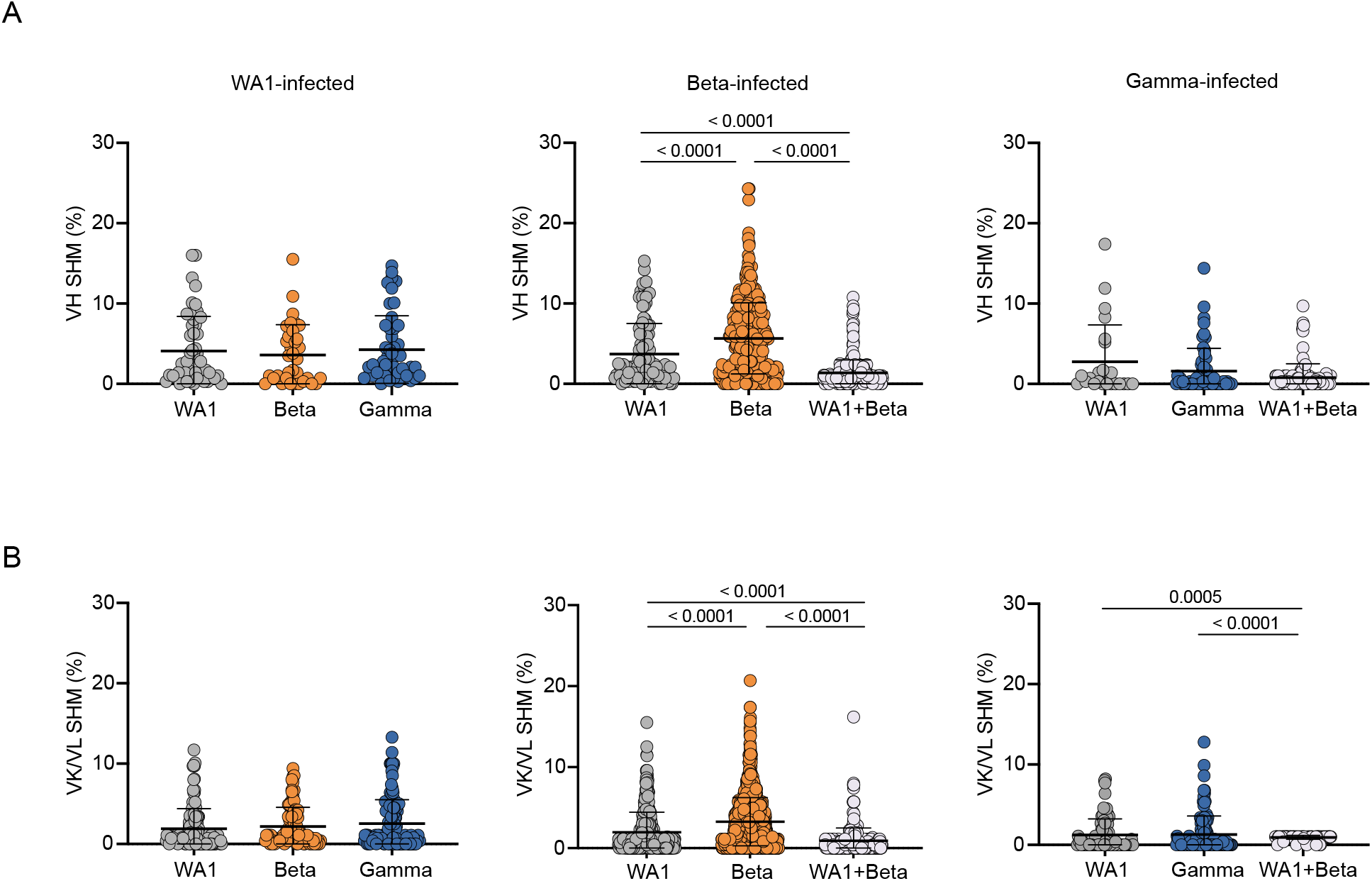
Somatic hypermutation (SHM) levels of SARS-CoV-2 specific B cells (unpaired sequences). SHM percent in variable heavy (V_H_) **(A)** or variable kappa/lambda (V_K_/V_L_) **(B)** regions. Error bars indicate the average number of nucleotide substitutions +/- standard deviation. Statistical significance was determined by the Mann-Whitney U test.

We next identified public clones in the SARS-CoV-2-specific repertoires elicited after infection with different variants. In total, 16 public clones were identified from 11 of the 13 infected individuals distributed across infection with all three variants (Fig. 5A). Notably, public clones for which data is available bound to Delta S-2P, and a subset of antibodies from the two most abundant public clones also bound to Omicron S-2P. One public clone, found in 5 individuals, uses IGHV4-59 with a short, strongly conserved, 6 amino acid CDR H3 and IGKV3-20 (Fig. 5, A-B). Antibodies matching the signature of public clone 1 have been previously described to bind the S2 domain of S and are generally cross-reactive with SARS-CoV-1^9, 20, 28^. Indeed, one member of this public clone was isolated from an individual infected with SARS-CoV-1^20^. This suggests that the convergent immune responses we observe may not be elicited only by variants of a single virus such as SARS-CoV-2 but can even extend to a broader range of related viruses.

**Fig. 5:**
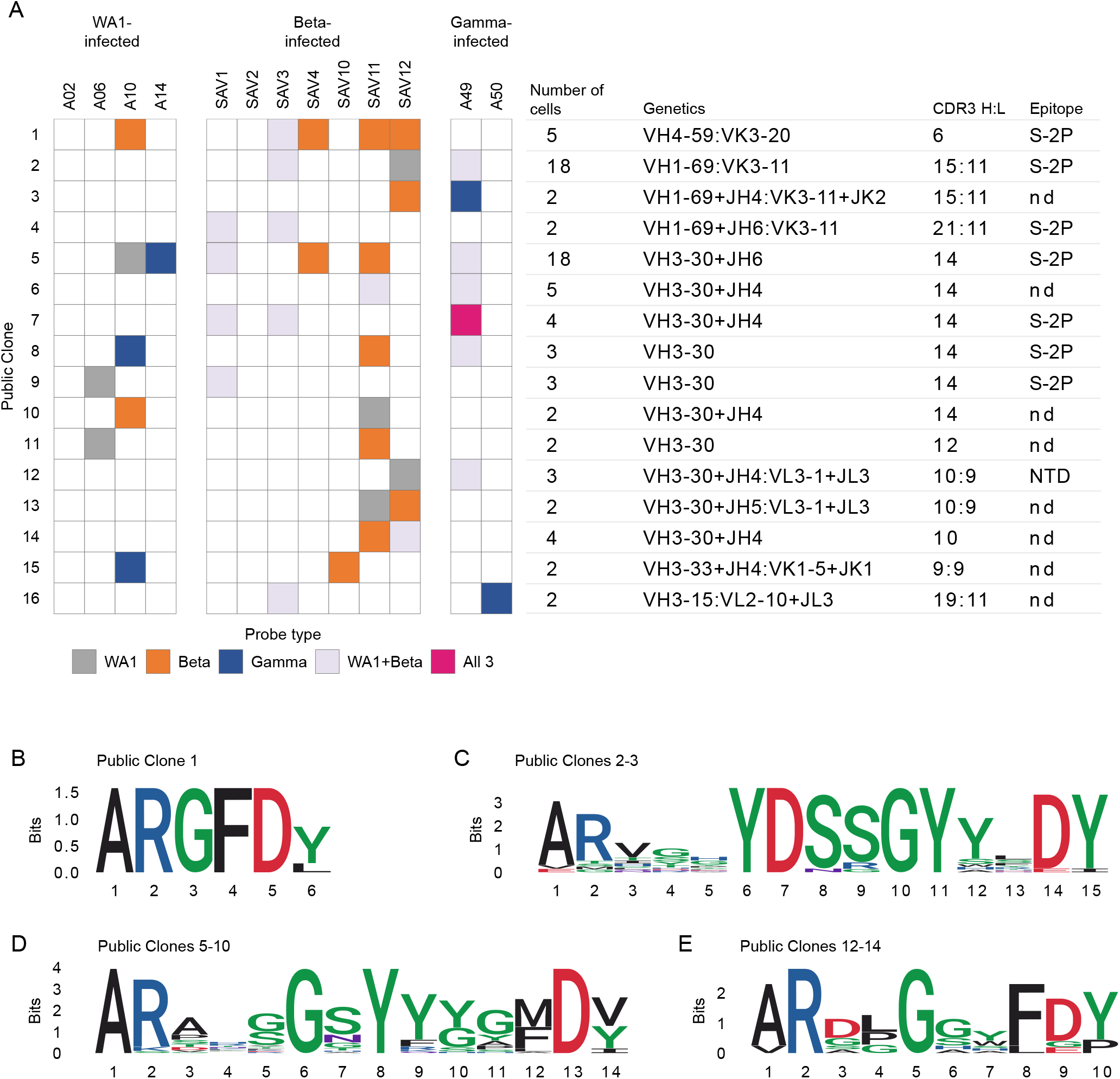
Public and cross-reactive clones. **(A)** Sixteen public clones were identified. Public clones are numbered 1-16 by row, as shown on the far left. Each column of boxes in the middle panel represents a single individual, as labeled at top, and is colored by probe(s) used, as shown at bottom. Right panel shows additional information about each public clone. Light chain information is provided after a colon if a consistent signature was found. Epitopes are inferred from ELISA of RATP-Ig supernatants of at least 1 public clone member; nd, not determined. **(B)** CDR H3 logogram for the top public clone, found in 5 of 13 individuals. (**C**)-(**E**) Combined CDR H3 logograms for **(C)** 2 public clones using IGHV1-69 and IGKV3-11 with a 15 amino acid CDR H3 length. **(D)** 6 public clones using IGHV3-30 with a 14 amino acid CDR H3 length. **(E)** 3 public clones using IGHV3-30 with a 10 amino acid CDR H3 length.

Public clones 2 and 3 both use the same heavy and light chain germline genes with the same CDR H3 and L3 lengths, though they fall outside of the 80% amino acid identity threshold. Combining sequences from both public clones revealed a strongly conserved IGHD3-22-encoded YDSSGY motif at positions 6-11 of CDR H3 (Fig. 5C). Strikingly, this is the same D gene implicated in targeting a Class IV RBD epitope^24^ although public clones 2 and 3 instead target an epitope in S2 and appear to be restricted to IGHV1-69 and IGKV3-11 V genes. We also observed the repeated use of IGHV3-30 with a 14 amino acid CDR H3 in six public clones which together comprise 35 cells from 8 different individuals. When we combined CDR H3 sequences from all 6 public clones in this group, we found an IGHD1-26-derived small-G-polar-Y-aromatic motif spanning positions 5-9 of CDR H3 (Fig. 5D). A large number of antibodies matching this signature have been previously described^7, 9, 20, 21, 22, 23, 28, 29^. The repeated observation of these closely related public clones in multiple individuals across all studied infection types further demonstrates the extraordinary convergence of the immune response to diverse variants.

We identified only one public clone, 12, that we were able to verify bound to either RBD or NTD, although public clones 13 and 14 also have highly similar V genes and CDR lengths (Fig. 5, A and E). Two previously reported antibodies, WRAIR-2038^30^ and COV-2307^22^, match the signature of these public clones and are also confirmed to bind NTD. The identification of a cross-reactive public clone is remarkable given deletions in Beta that disrupt the main NTD supersite for neutralizing antibodies^31^. This again highlights the capacity of the adaptive immune response to find consistent ways to target the SARS-CoV-2 virus, despite substitutions selected for their ability to disguise the targets.

A deep understanding of the IG repertoires that mediate cross-protective responses to SARS-CoV-2 after infection or vaccination will be critical for guiding therapeutic approaches to future variants as the virus continues to evolve. In this study, we used rapid mAb production and functional analysis, and single cell Ig sequencing to conduct an in-depth, unbiased characterization of total antigen-specific B cell responses against multiple SARS-CoV-2 variants including Delta and Omicron from people infected with the ancestral WA1, Beta, or Gamma. Our principal findings were: 1) infection with any of the “older” variants consistently elicited substantial numbers of antibodies capable of cross-binding even to the more recent antigenically divergent variants Delta and Omicron; 2) infection with any of these variants elicited antibodies targeting the same immunodominant epitopes in RBD; 3) antigen-specific memory B cells elicited by SARS-CoV-2 are polyclonal and use similar patterns of heavy and light chain V genes, irrespective of the infecting variant; and 4) public clones and other cross-reactive antibodies are common among responses to all infecting variants. Our results demonstrate a fundamentally convergent humoral immune response across different SARS-CoV-2 variants that cross-bind even to antigenically distant ones such as Delta and Omicron.

To date, most analyses of SARS-CoV-2-specific B cells have focused on neutralizing antibodies with potential therapeutic applications. Those which have investigated the total binding repertoire have used samples from people infected with the ancestral WA1 variant^7, 10^; here we extend such analysis to individuals infected with the antigenically distinct Beta and Gamma variants and show that antibodies capable of binding to multiple variants are common. Indeed, while the strength of cross-neutralization depends on the antigenic distance from the infecting variant^32^, we found that most WA1-Beta cross-binding antibodies can also bind to a later, more divergent, variants such as Delta, and approximately half can additionally bind Omicron.

Furthermore, we observed that the hierarchy of immunodominant epitopes targeted on these variants remains unchanged. While a recent report found that Beta-infection was less likely to elicit antibodies contacting S residue F456 than WA1-infection^33^, we found no changes in targeting of the RBD-A epitope, which includes this residue. Interestingly, even though the immunodominance of binding epitopes is known to be consistent in response to WA1, Beta, or Omicron mRNA immunization^18, 19^, recent reports have found that infection with an Omicron subvariant after vaccination can shift the epitope landscape compared to vaccination alone^34, 35^. This likely reflects the effect of imprinting by consecutive exposures to closely related antigens^36^, although differences in the primary response to Omicron variants cannot yet be ruled out. For earlier variants, at least, we demonstrate here similar patterns of immunodominance after variant infection, a phenomenon that may help explain the continued efficacy of vaccines based on ancestral variants.

In addition to concordant epitope targeting, we also found consistent IG V gene usage in the antibody response to all three variants we investigated. Our findings highlight the difference between the neutralizing antibody repertoires investigated previously compared to the total binding repertoires examined here, emphasizing the insights to be gleaned by taking a broader perspective. Thus, while many of the variant-induced public clones that were cross-reactive with all three variants, as well as Delta and sometimes Omicron, appear to be non-neutralizing and S2 domain-binding, the breadth and ready elicitation may be important for Fc-dependent functions ^30, 37^. Therefore, public clones stimulated by one variant could play a protective role against later variants, even when neutralizing antibodies are less effective. Overall, more than 8% of the cells that we sequenced belong to a public clone, highlighting again the extraordinary convergence of the antibody response across antigenically distinct variants of SARS-CoV-2. Importantly, even when sequence homology fell below the threshold to define clones as public, we found conserved motifs which are likely to drive functional convergence consistent with recent evidence that antibodies may target overlapping epitopes using comparable binding conformations in the absence of convergent V genes^38^. Together, these findings further highlight the capability of the human immune system to respond to SARS-CoV-2 in a manner that is largely conserved yet at the same time tolerant of differences between variants.

In summary, our data reveal marked convergence that defines multiple aspects of the humoral immune response to different SARS-CoV-2 variants. This phenomenon comprises convergent V-gene usage and epitope specificities elicited by primary exposure to SARS-CoV-2 variants, including a substantial proportion of public clones and cross-binding B cells. This suggests the existence of immunological constraints guiding the response to related viruses, even in the face of substantial antigenic divergence, and may explain how first-generation vaccine designs using the ancestral S protein sequence have generally proven equally as protective against severe disease compared to updated vaccines matched to recent variants^39, 40^.

Our study is limited by sampling of paired heavy and light chain sequences from fewer than 1,000 SARS-CoV-2-specific B cells across 13 individuals. This scale is small in comparison to bulk IG sequencing studies and even a few single-cell studies. We are also limited in our ability to make functional repertoire comparisons due to varied sorting strategies and differences in functional assays used to assess isolated mAbs. Moreover, our cohort was sampled only at a single time point early in convalescence and included only one individual with high serum neutralization titers. It will be important to verify that our findings extend to later time points when the antibody repertoire has matured. In addition, while Beta and Gamma are antigenically distinct from WA1, they only represent a small portion of the SARS-CoV-2 antigenic map^41^. Further studies are needed to examine the response elicited by more antigenically divergent SARS-CoV-2 variants such as Delta and Omicron.

## MATERIALS AND METHODS

### Study design

We selected 13 convalescent individuals that had experienced symptomatic Covid-19 infection with either WA1 virus or the Beta or Gamma variants. Serum, plasma and PBMC were isolated at each respective clinical center. The selection of individuals was based on the availability of samples collected at similar time-points (between 17 and 38 days after symptoms onset), rather than the severity of disease or neutralizing antibody titers (Extended Data Fig. 1). Seven individuals were infected with the Beta variant and recruited at the Sheba Medical Center, Tel HaShomer, Israel. Because of limited sample availability, two additional Beta-infected individuals were recruited at the Vaccine Research Center (VRC) and used for T cell analyses. Two individuals were infected with the Gamma variant and recruited at the University of Minnesota Hospital, USA. Infections with Beta and Gamma variants were confirmed by sequencing. The samples from four WA1-infected individuals, collected early in the pandemic prior to the emergence of variants, as well as the two additional beta-infected individuals used for T cell analysis were collected under the VRC, National Institute of Allergy and Infectious Diseases (NIAID), National Institutes of Health’s protocol VRC 200 (NCT00067054) in compliance with the NIH Institutional Review Board (IRB) approved protocol and procedures. All subjects met protocol eligibility criteria and agreed to participate in the study by signing the NIH IRB approved informed consent. Research studies with these samples were conducted by protecting the rights and privacy of the study participants. All participants provided informed consent in accordance with protocols approved by the respective institutional review boards and the Helsinki Declaration.

### Serology

Antibody binding was measured by 10-plex Meso Scale Discovery Electrochemiluminescence immunoassay (MSD-ECLIA) as previously described^4^. Cell-surface S binding was assessed as previously described^4^. Serum neutralization titers for either WA1-D614G, Beta, Gamma or Delta pseudotyped virus particles were obtained as previously described^4^.

### Antigen-specific ELISA

Reacti-Bind 96-well polystyrene plates (Pierce) were coated with 100 μl of affinity purified goat anti-human IgG Fc (Rockland) at 1:20,000 in PBS, or 2 μg/ml SARS-CoV-2 recombinant protein in PBS overnight at 4°C. Plates were washed in PBS-T (500ml 10XPBS + 0.05% Tween-20 + 4.5L H2O) and blocked for 1 h at 37°C with 200 μL/well of B3T buffer: 8.8 g/liter NaCl, 7.87 g/liter Tris-HCl, 334.7 mg/liter EDTA, 20 g BSA Fraction V, 33.3 ml/liter fetal calf serum, 666 ml/liter Tween-20, and 0.02% Thimerosal, pH 7.4). Diluted antibody samples were applied and incubated 1 hr at 37°C followed by 6 washes with PBS-T; plates were the incubated with HRP-conjugated anti-human IgG (Jackson ImmunoResearch) diluted 1:10,000 in B3T buffer for 1 h at 37°C. After 6 washes with PBS-T, SureBlue TMB Substrate (KPL) was added, incubated for 10 min, and the reaction was stopped with 1N H2SO4 before measuring optical densities at 450nm (Molecular Devices, SpectraMax using SoftMax Pro 5 software). For single-point assays, supernatants from transfected cells were diluted 1:10 in B3T and added to the blocked plates. Purified monoclonal antibodies were assessed using 5-fold serial dilutions starting at 10ug/ml. To assess the levels of IgG in supernatants, standard curves were run on the same plates as supernatants, using threefold serial dilutions of human IgG (Sigma) starting at 100ng/ml IgG. ELISA signals were considered positive if they were greater than or equal to 2X the average of the blank wells of the plate.

### Pseudovirus neutralization assay

SARS-CoV-2 spike pseudotyped lentiviruses were produced by co-transfection of 293T cells with plasmids encoding the lentiviral packaging and luciferase reporter, a human transmembrane protease serine 2 (TMPRSS2), and SARS-CoV-2 S genes using Lipofectamine 3000 transfection reagent (ThermoFisher, CA)^15, 42^. Forty-eight hours after transfection, supernatants containing pseudoviral particles were harvested, filtered and frozen. For neutralization assay two dilutions of the transfection supernatants (2- or 3-fold) were mixed with equal volume of titrated pseudovirus (final dilution 4x or 6x), incubated for 45 minutes at 37 °C and added to pre-seeded 293 flpin-TMPRSS2-ACE2 cells (made by Adrian Creanga, VRC, NIH) in triplicate in 96-well white/black Isoplates (Perkin Elmer). Following 2 hours of incubation, wells were replenished with 150 µL of fresh medium. Cells were lysed 72 hours later and luciferase activity (relative light unit, RLU) was measured. Percent neutralization was calculated relative to pseudovirus-only wells.

### Intracellular cytokine staining

The T cell staining panel used in this study was modified from a panel developed by the laboratory of Dr. Steven De Rosa (Fred Hutchinson Cancer Research Center). Directly conjugated antibodies purchased from BD Biosciences include CD19 PE-Cy5 (Clone HIB19; cat. 302210), CD14 BB660 (Clone M0P9; cat. 624925), CD3 BUV395 (Clone UCHT1; cat. 563546), CD4 BV480 (Clone SK3; cat. 566104), CD8a BUV805 (Clone SK1; cat. 612889), CD45RA BUV496 (Clone H100; cat. 750258), CD154 PE (Clone TRAP1; cat. 555700), IFNg V450 (Clone B27; cat. 560371 and IL-2 BB700 (Clone MQ1-17H12; cat. 566404). Antibodies from Biolegend include CD16 BV570 (Clone 3G8; cat. 302036), CD56 BV750 (Clone 5.1H11; cat. 362556), CCR7 BV605 (Clone G043H7; cat. 353244) and CD69 APC-Fire750 (Clone FN50; cat. 310946). TNF FITC (Clone Mab11; cat. 11-7349-82) and the LIVE/DEAD Fixable Blue Dead Cell Stain (cat. L34962) were purchased from Invitrogen.

Cryopreserved PBMC were thawed into pre-warmed R10 media (RPMI 1640, 10% FBS, 2 mM L-glutamine, 100 U/ml penicillin, and 100 μg/ml streptomycin) containing DNase and rested for 1 hour at 37°C/5% CO_2_. For stimulation, 1 – 1.5 million cells were plated into 96-well V-bottom plates in 200mL R10 and stimulated with SARS-CoV-2 peptide pools (2ug/mL for each peptide) in the presence of Brefeldin A (Sigma-Aldrich) and monensin (GolgiStop; BD Biosciences) for 6 hours at 37°C/5%CO_2_. A DMSO-only condition was used to determine background responses. Following stimulation samples were stained with LIVE/DEAD Fixable Blue Dead Cell Stain for 10 minutes at room temperature and surface stained with titrated amounts of anti-CD19, anti-CD14, anti-CD16, anti-CD56, anti-CD4, anti-CD8, anti-CCR7 and anti-CD45RA for 20 minutes at room temperature. Cells were washed in FACS Buffer (PBS + 2% FBS), and fixed and permeabilized (Cytofix/Cytoperm, BD Biosciences) for 20 minutes at room temperature. Following fixation, cells were washed with Perm/Wash buffer (BD Biosciences) and stained intracellularly with anti-CD3, anti-CD154, anti-CD69, anti-IFNg, anti-IL-2 and anti-TNF for 20 minutes at room temperature. Cells were subsequently washed with Perm/Wash buffer and fixed with 1% paraformaldehyde. Data were acquired on a modified BD FACSymphony and analyzed using FlowJo software (version 10.7.1). Cytokine frequencies were background subtracted and negative values were set to zero.

Synthetic peptides (>75% purity by HPLC; 15 amino acids in length overlapping by 11 amino acids) were synthesized by GenScript. To measure T cell responses to the full-length WA-1 S glycoprotein (YP_009724390.1), 2 peptide pools were utilized, S pool A (peptides 1-160; residues 1-651) and S pool B (peptides 161-316; residues 641-1273) (Supplementary Table 4). Peptides were 15 amino acids in length and overlapped by 11 amino acids. S pool A contained peptides for both D614 and the G614 mutation. Responses to full-length S were calculated by summing the responses to both pools after background subtraction. Select peptide pools were used to measure T cell responses to mutated regions of the S glycoproteins of the Alpha, Beta and Gamma SARS-CoV-2 variants along with control pools corresponding to the same regions within the WA-1 S glycoprotein (Supplementary Table 5).

### Epitope mapping by Surface Plasmon Resonance (SPR)

Serum epitope mapping competition assays were performed, as previously described ^18, 19^, using the Biacore 8K+ surface plasmon resonance system (Cytiva). Anti-histidine antibody was immobilized on Series S Sensor Chip CM5 (Cytiva) through primary amine coupling using a His capture kit (Cytiva). Following this, his-tagged SARS-CoV-2 S protein containing 2 proline stabilization mutations (S-2P) was captured on the active sensor surface.

Human IgG monoclonal antibodies (mAb) used for these analyses include: B1-182, CB6, A20-29.1, A19-46.1, LY-COV555, A19-61.1, S309, A23-97.1, A19-30.1, A23-80.1, and CR3022. Either competitor or negative control mAb was injected over both active and reference surfaces. Human sera were then flowed over both active and reference sensor surfaces, at a dilution of 1:50. Following the association phase, active and reference sensor surfaces were regenerated between each analysis cycle.

Prior to analysis, sensorgrams were aligned to Y (Response Units) = 0, using Biacore 8K Insights Evaluation Software (Cytiva), at the beginning of the serum association phase. Relative “analyte binding late” report points (RU) were collected and used to calculate percent competition (% C) using the following formula: % C = [1 – (100 * ((RU in presence of competitor mAb) / (RU in presence of negative control mAb))]. Results are reported as percent competition and statistical analysis was performed using unpaired, two-tailed t-test (Graphpad Prism v.8.3.1). All assays were performed in duplicate and averaged.

Only one of the WA1-infected individuals (A14) produced sufficiently high binding titers against Beta and Delta S to enable epitope mapping by competition. In addition, Beta-infected donors SAV2 and SAV10 were below the lower limit of quantification for WA1 and Delta S.

### Production of antigen-specific probes

Biotinylated probes for S-2P, NTD and RBD were produced as described previously ^43, 44^. Briefly, single-chain Fc and AVI-tagged proteins were expressed transiently for 6 days. After harvest, the soluble proteins were purified and biotinylated in a single protein A column followed by final purification on a Superdex 200 16/600 gel filtration column. Biotinylated proteins were then conjugated to fluorescent streptavidin.

### Antigen-specific B cell sorting

PBMC vials containing approximately 10^7^ cells were thawed and stained with Live/Dead Fixable Blue Dead Cell Stain Kit (Invitrogen, cat# L23105) for 10 min at room temperature, followed by incubation for 20 min with the staining cocktail consisting of antibodies and probes. The antibodies used in the staining cocktail were: CD8-BV510 (Biolegend, clone RPA-T8, cat# 301048), CD56-BV510 (Biolegend, clone HCD56, cat# 318340), CD14-BV510 (Biolegend, clone M5E2, cat# 301842), CD16-BUV496 (BD Biosciences, clone 3G8, cat# 612944), CD3-APC-Cy7 (BD Biosciences, clone SP34-2, cat# 557757), CD19-PECy7 (Beckmann Coulter, clone J3-119, cat# IM36284), CD20 (BD Biosciences, clone 2H7, cat# 564917), IgG-FITC (BD Biosciences, clone G18-145, cat# 555786), IgA-FITC (Miltenyi Biotech, clone IS11-8E10, cat# 130-114-001) and IgM-PECF594 (BD Biosciences, clone G20-127, cat# 562539). For each variant, a set of two S probes S-2P-APC and S-2P-BUV737, in addition to RBD-BV421 and NTD-BV711 were included in the staining cocktail for flow cytometry sorting.

For RATP-Ig, single-cells were sorted in 96-well plates containing 5 µL of TCL buffer (Qiagen) with 1% β-mercaptoethanol according to the gating strategy shown in Fig. S2B. Samples sorted for 10x Genomics single-cell RNAseq were individually labelled with an oligonucleotide-linked hashing antibody (Totalseq-C, Biolegend) in addition to the staining cocktail and sorted into a single tube according to the gating strategy shown in Fig. S2B. All cell sorts were performed using a BD FACSAria II instrument (BD Biosciences). Frequency of antigen-specific B cells were analyzed using FlowJo 10.8.1 (BD Biosciences).

### Monoclonal antibody isolation and characterization by RATP-Ig

#### cDNA synthesis

Variable heavy and light chains were synthesized using a modified SMARTSeq-V4 protocol by 5’ RACE. Single-cell RNA was first purified with RNAclean beads (Beckman Coulter). cDNA was then synthesized using 5’ RACE reverse-transcription, adding distinct 3’ and 5’ template switch oligo adapters to total cDNA. cDNA was subsequently amplified with TSO_FWD and TS_Oligo_2_REV primers. Excess oligos and dNTPs were removed from amplified cDNA with EXO-CIP cleanup kit (New England BioLabs).

#### Immunoglobulin enrichment and sequencing

Heavy and light chain variable regions were enriched by amplifying cDNA with TSO_FWD and IgA/IgG_REV or IgK/IgL_REV primer pools. An aliquot of enriched product was used to prepare Nextera libraries with Unique Dual Indices (Illumina) and sequenced using 2×150 paired-end reads on an Illumina MiSeq. Separate aliquots were used for IG production; RATP-Ig is a modular system and can produce single combined or separate HC/LC cassettes.

#### Cassette fragment synthesis

Final cassettes include CMV, and HC/LC-TBGH polyA fragments. To isolate these fragments, amplicons were first synthesized by PCR. PCR products were run on a 1% agarose gel and fragments of the correct length were extracted with Thermo gel extraction and PCR cleanup kit (ThermoFisher Scientific). Gel-extracted products were digested with DpnI (New England Biolabs) to further remove any possible contaminating plasmid. These fragment templates were then further amplified to create final stocks of cassette fragments.

#### Cassette assembly

Enriched variable regions were assembled into linear expression cassettes in two sequential ligation reactions. The first reaction assembles CMV-TSO, TSO-V-LC, and KC-IRES fragments into part 1 and IRES-TSO, TSO-V-HC, and IgGC-TBGH fragments into part 2 using NEBuilder HIFI DNA Assembly Mastermix (New England BioLabs). Following reaction 1, parts 1 and 2 were combined into a single reaction 2 and ligated into a single cassette.

Separate cassettes: Enriched variable regions were assembled into linear expression cassettes by ligating CMV-TSO, TSO-V-C, and C-TBGH fragments using NEBuilder HIFI DNA Assembly Mastermix (New England BioLabs). Assembled cassettes were amplified using CMV_FWD and TBGH_REV primers. Amplified linear DNA cassettes encoding monoclonal heavy and light chain genes were co-transfected into Expi293 cells in 96-well deep-well plates using the Expi293 Transfection Kit (ThermoFisher Scientific) according to the manufacturer’s protocol. Microtiter cultures were incubated at 37 degrees and 8% CO_2_ with shaking at 1100 RPM for 5-7 days before supernatants were clarified by centrifugation and harvested. It is important to note that supernatant IgG titers were not calculated but were only verified to reach a minimum cutoff value for functional assays, limiting our ability to compare potency between antibodies.

### Droplet-based single cell isolation and sequencing

Antigen-specific memory B cells were sorted as described above. Cells from two separate sorts were pooled in a single suspension and loaded on the 10x Genomics Chromium instrument with reagents from the Next GEM Single Cell 5’ Kit v1.1 following the manufacturer’s protocol to generate total cDNA. Heavy and light chains were amplified from the cDNA using custom 3’ primers specific for IgG, IgA, IgK or IgL with the addition of Illumina sequences^45^. The Illumina-ready libraries were sequenced using 2×300 paired-end reads on an Illumina MiSeq. Hashing oligonucleotides were amplified and sequenced from the total cDNA according to the 10x Genomics protocol.

### V(D)J sequence analysis

For cells processed via RATP-Ig, reads were demultiplexed using a custom script and candidate V(D)J sequences were generated using BALDR^46^ and filtered for quality using a custom script. The resulting sequences were annotated using SONAR v4.2^47^ in single-cell mode.

For cells processed via the 10x Genomics Chromium device, reads from the hashing libraries were processed using cellranger (10x Genomics). The resulting count matrix was imported into Seurat^48^ and the sample of origin called using the HTODemux function. Paired-end reads from V(D)J libraries were merged and annotated using SONAR in single-cell mode with UMI detection and processing.

For all datasets, nonproductive rearrangements were discarded, as were any cells with more than one productive heavy or light chain. Cells with an unpaired heavy or light chain were included in calculations of SHM and gene usage statistics, but were excluded from assessments of clonality and determination of public clones. Public clones were determined by using the clusterfast algorithm in vsearch^49^ to cluster CDR H3 amino acid sequences at 80% identity. Where relevant, all clonally related B cells in a single individual were included in a public clone, even if not all were directly clustered together in the vsearch analysis. While light chain V genes and CDR3 were not used to define public clones, they are reported when we found a consistent signature within a public clone.

## Acknowledgements

The authors would like to thank the members of the VRC 200 Study Team for their role in collecting samples that were used in this study: Lesia Drupolic, Lasonji Holman, Maria Burgos Florez, Charla Andrews, Britta Flach, Emily Coates, Obrimpong Amoa-Awua, Jennifer Cunningham, Pamela Costner, Floreliz Mendoza, William Whalen, Jamie Saunders, Laura Novik, Aba Eshun, Anita Arthur, Xiaolin Wang, Karen Parker, Abidemi Ola, Catina Evans, Jennifer Phipps, Pernell Williams, Justine Jones, Jackie Stephens, Jumoke Gbadebo, Preeti Apete, Renunda Hicks, LaShawn Requillman, Alison Beck, Seemal Awan, Richard Wu, Priya Kamath, Olga Trofymenko, Sarah Plummer, Nina Berkowitz, Olga Vasilenko, and Iris Pittman. The authors also thank Dr. Steven De Rosa (Fred Hutchinson Cancer Center) for providing a 28-color flow cytometry panel which we modified for our study and David Ambrozak for assistance with cell sorting.

The authors thank Rodrigo Matus Nicodemus for his assistance in developing primers for RATP-Ig.

## Author Contributions

Conceptualization: NSL, CAS, DCD

Data curation: MM, CAS

Formal Analysis: NSL, MM, TSJ, DAW, LW, KB, SRN, SOC, KLB, CAS

Investigation: NSL, MM, TSJ, DAW, ARH, LW, KB, WPB, SDS, DM, CGL, BZ, KLB, JRT, RLD, LP, JW, CAT

Methodology: TSJ, DCD

Resources: ESY, YZ, SOD, MC, AS, KL, WS, RK, AB, TZ, JR, SV, AA, LN, AW, IG, M Guech, ITT, EP, TJR

Supervision: AP, JM, NADR, M Guadinski, RAK, PDK, ABM, SA, TWS, IL, JRM, NJS, CAS, DCD

Visualization: NSL, MM, TSJ, DAW, KLB, CAS, DCD

Writing – original draft: NSL, MM, TSJ, CAS, DCD

Writing – review & editing: all authors

## Data and materials availability

All data and materials are available upon request.

## Extended Data

**Fig. 1:**
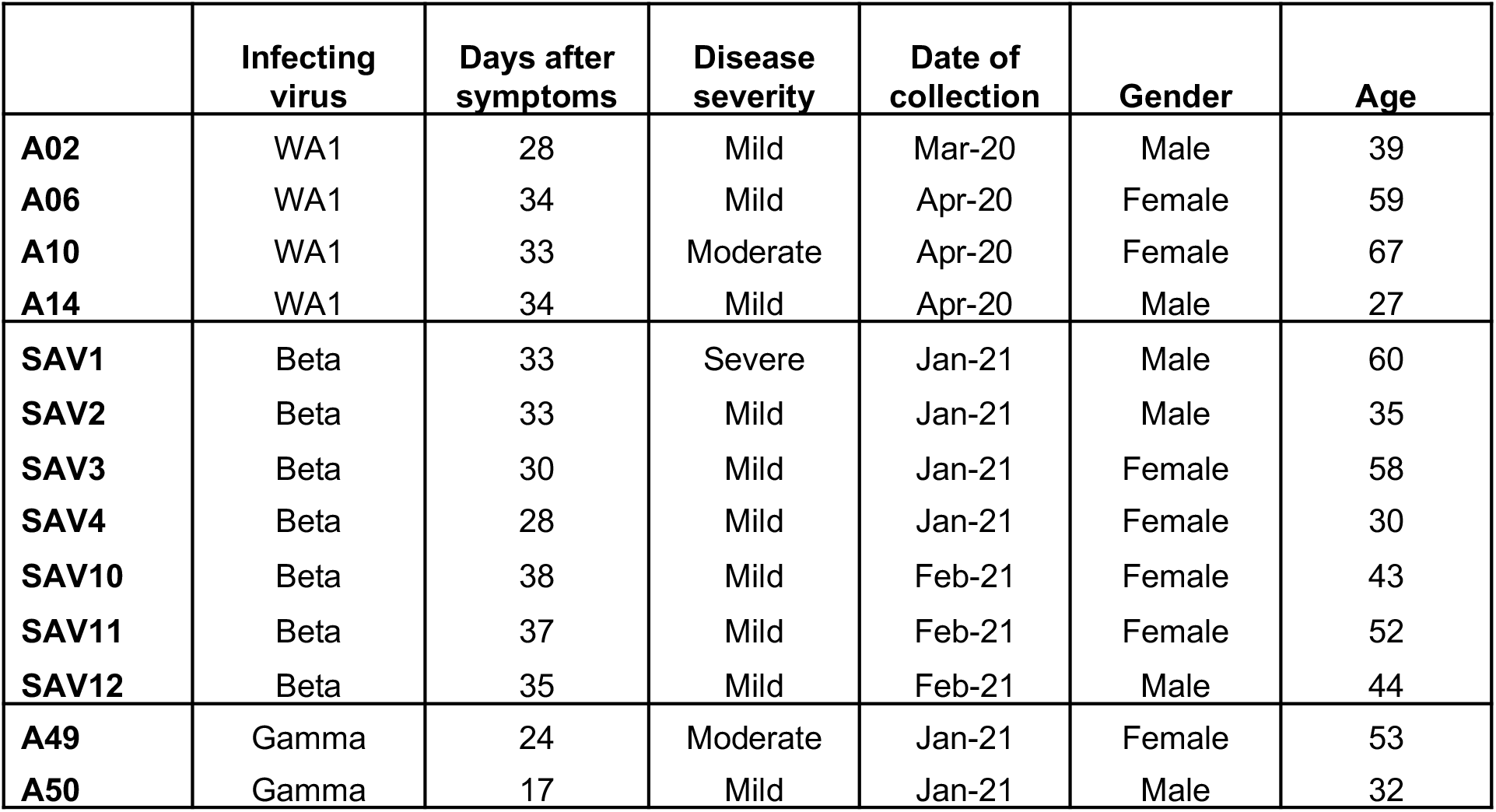
Details of the study cohort.

**Fig. 2:**
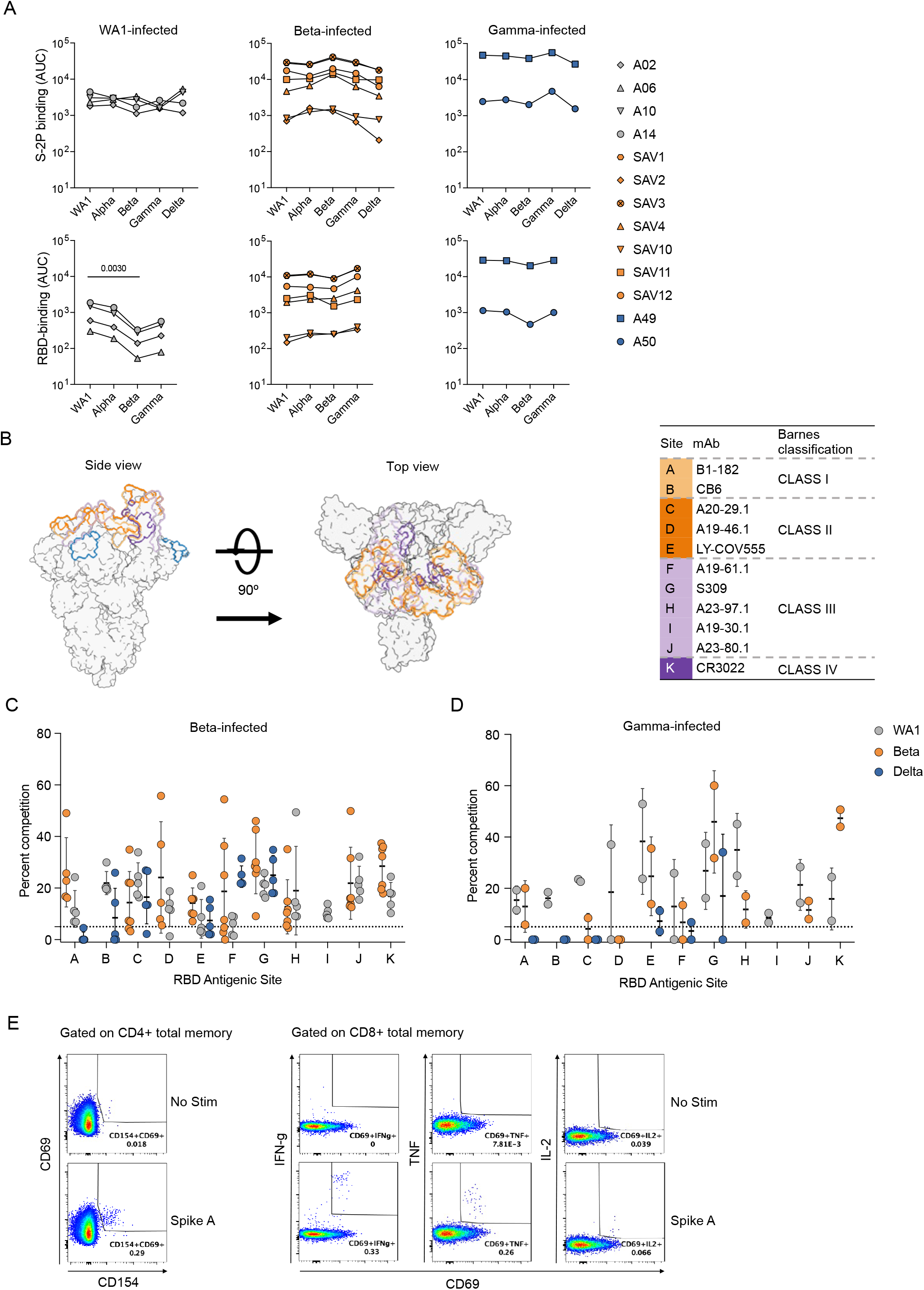
Additional serology and epitope mapping data. **(A)** Binding antibody titers to spike (top panels) and RBD (bottom panels) from different variants indicated on the x-axis. **(B)** Structural schematic of spike protein showing epitopes from monoclonal antibodies used for RBD epitope mapping by competition assay; **(C)** Epitope mapping of Beta-infected individuals on WA1, Beta and Delta spike proteins; **(D)** Epitope mapping of Gamma-infected individuals on WA1, Beta and Delta spike proteins; **(E)** Gating strategy for T cell response analysis.

**Fig. 3:**
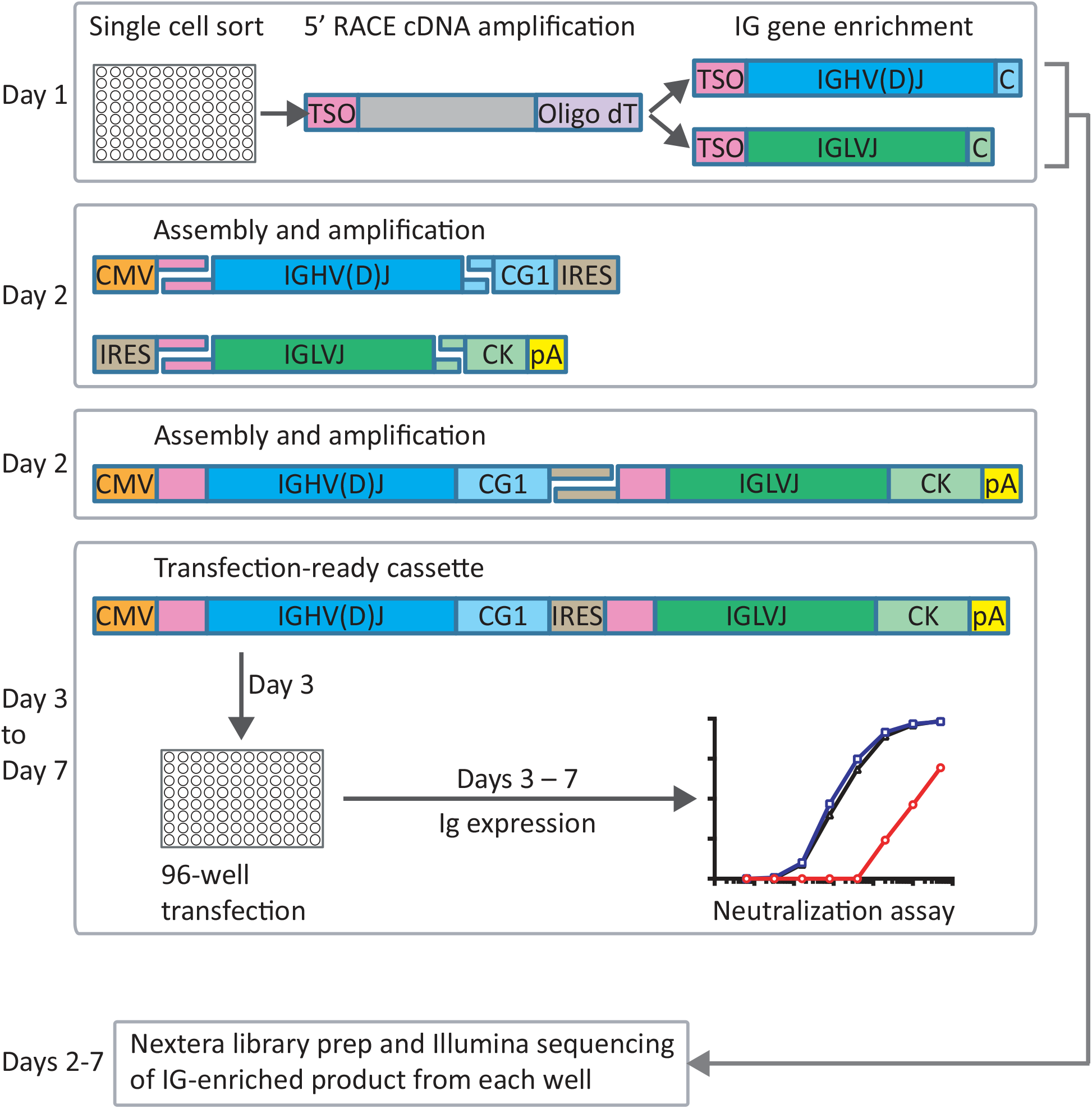
Rapid assembly, transfection and production of immunoglobulin (RATP-Ig) workflow. 5’-RACE is used to generate total cDNA. Full-length heavy and light chain immunoglobulin V genes are enriched by PCR and assembled into recombinant mAb linear expression cassettes. In parallel, V gene libraries are synthesized and sequenced by NGS. Final cassettes are transfected into 96-well Expi293 microtiter cultures, and culture supernatants are collected up to 7 days after initial sort for functional screening.

**Fig. 4:**
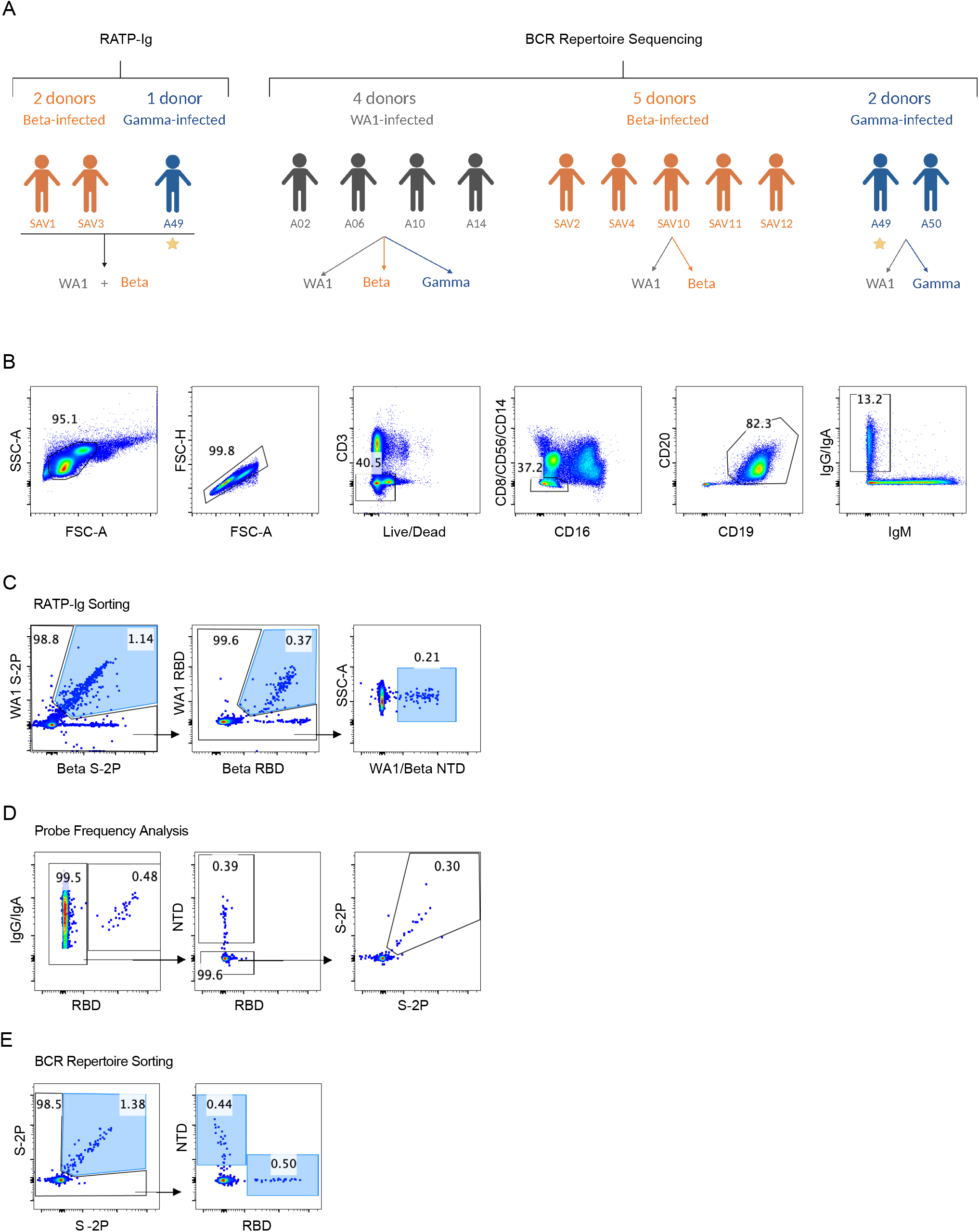
Antigen-specific B cell sorting. (**A**) Arrows indicate probes used for sorting antigen-specific B cells from each group of convalescent individuals. The individual marked with a star was used for both RATP-Ig and total BCR repertoire sequencing. (**B**) Flow cytometry representative plots and gating strategy for class-switched memory B cells. **(C)-(E)** Representative plots and gating strategy for sorting and analysis of antigen-specific cells for **(C)** RATP-Ig, **(D)** Frequency analysis, and **(E)** Repertoire sequencing. Final sort gates are shown in blue.

**Fig. 5:**
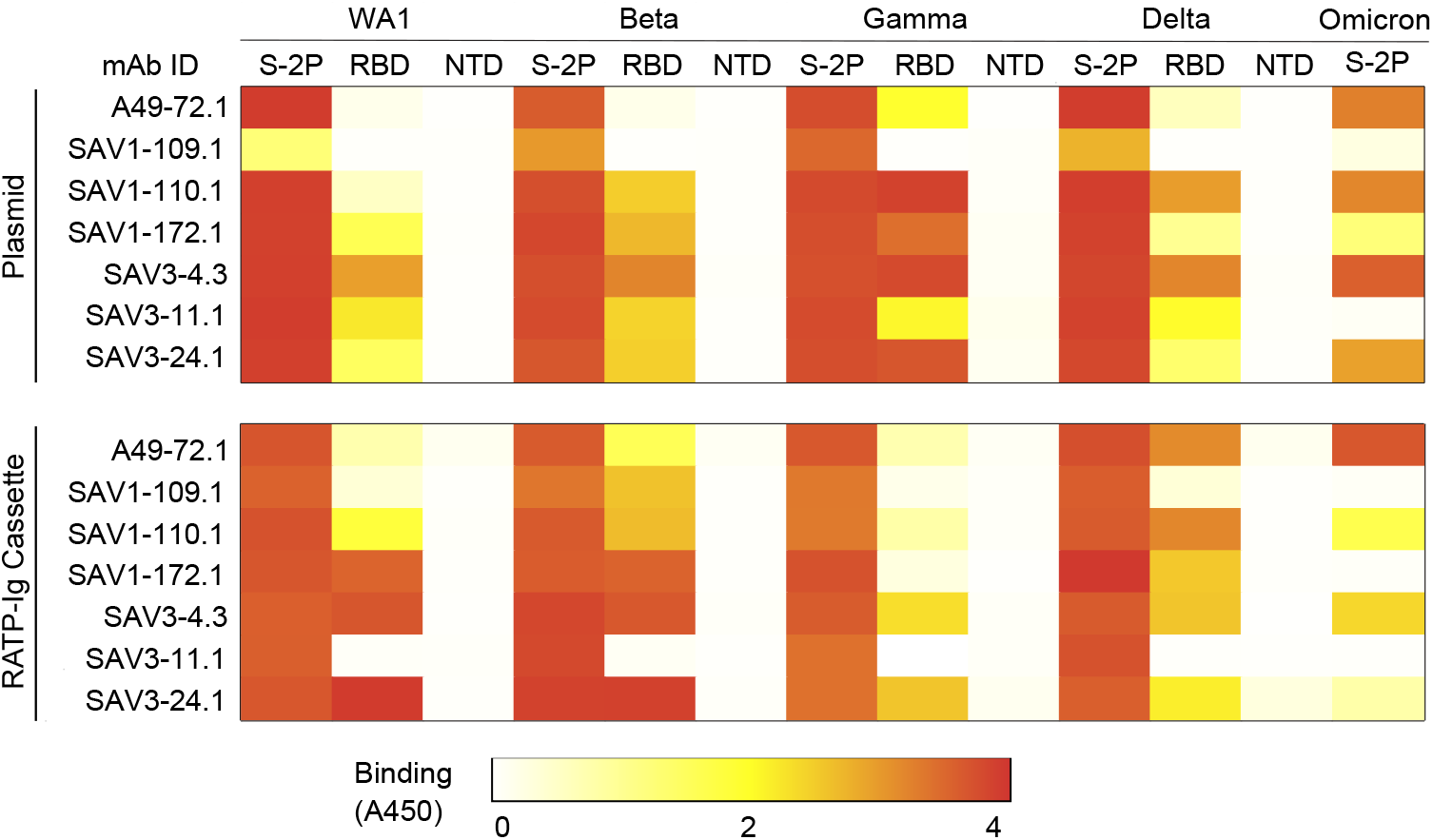
Validation of RATP-Ig screening with synthesized plasmids. Heatmaps show ELISA absorbance at 450 nm (not quantitative).

**Fig. 6:**
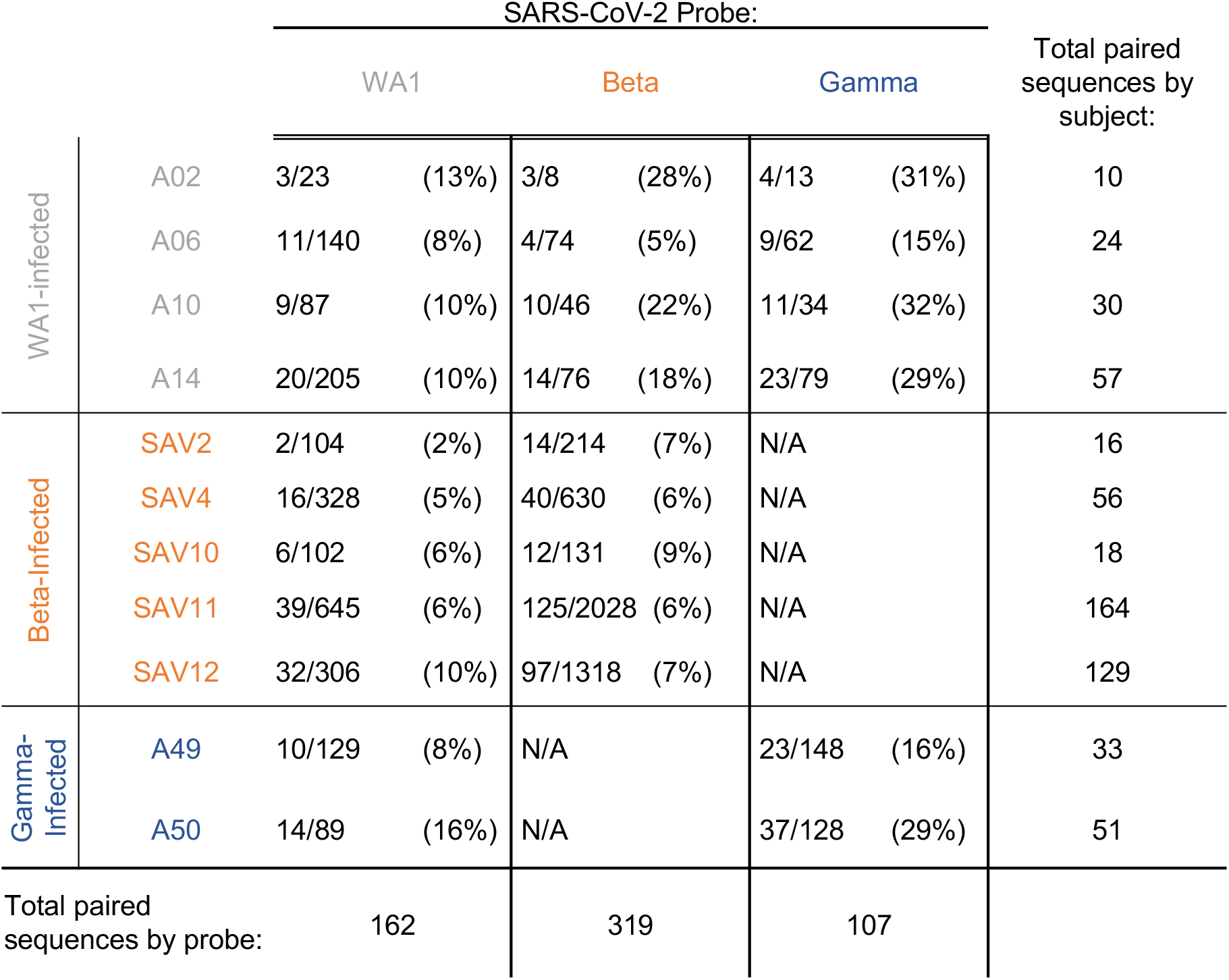
Sample recovery from 10x Genomics-based single cell isolation and sequencing.

**Fig. 7:**
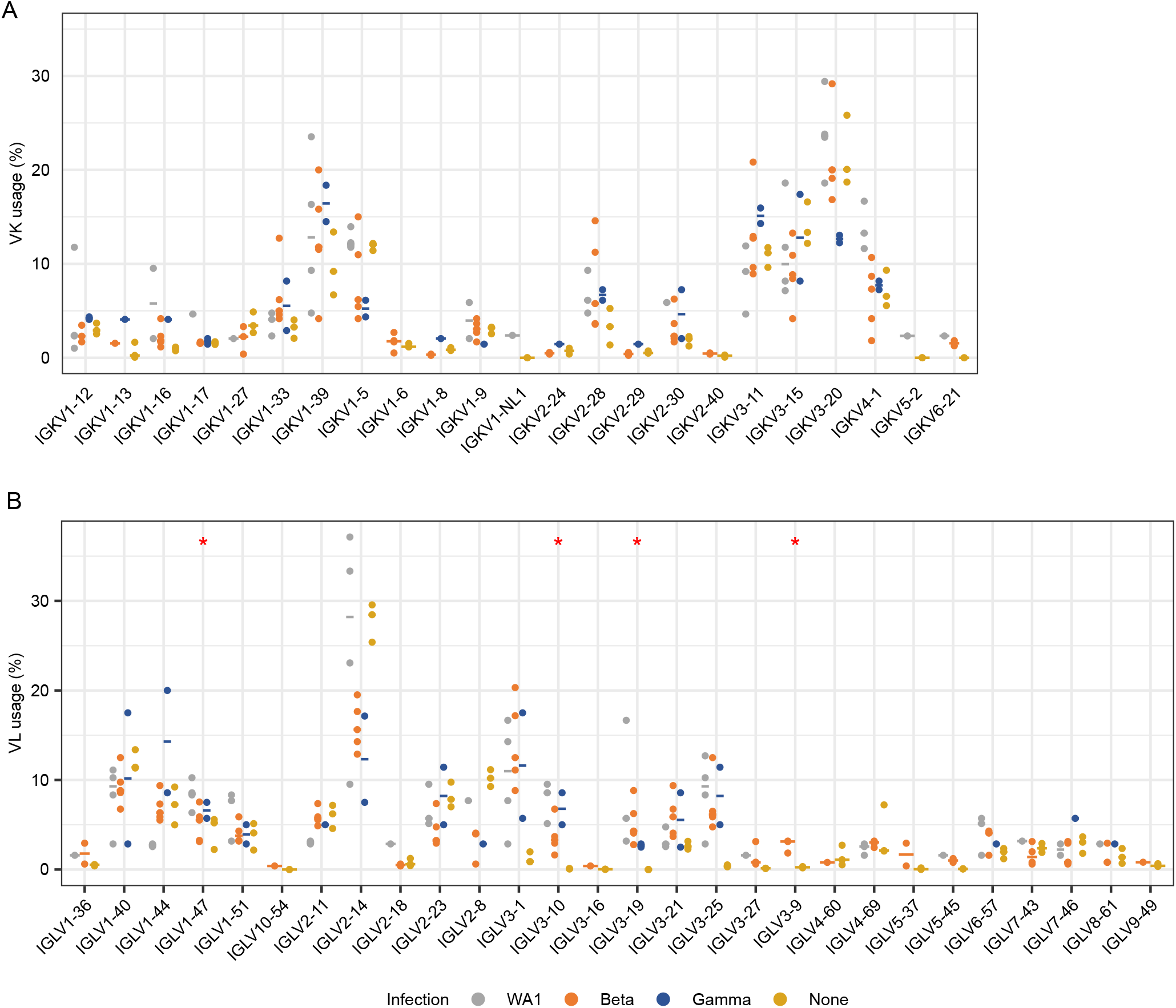
SARS-CoV-2-specific light chain V gene usage frequencies. **(A)** Kappa and **(B)** Lambda chain V gene repertoire analysis by infecting variant, with WA1, Beta and Gamma shown in grey, orange and blue, respectively, and data from pre-pandemic controls in yellow. The x-axis shows all germline genes used; the y-axis represents the percent of individual gene usage. Stars indicate genes with at least one significant difference between groups; pairwise comparisons using the Dunn test are in Extended Data Fig. 8.

**Fig. 8:**
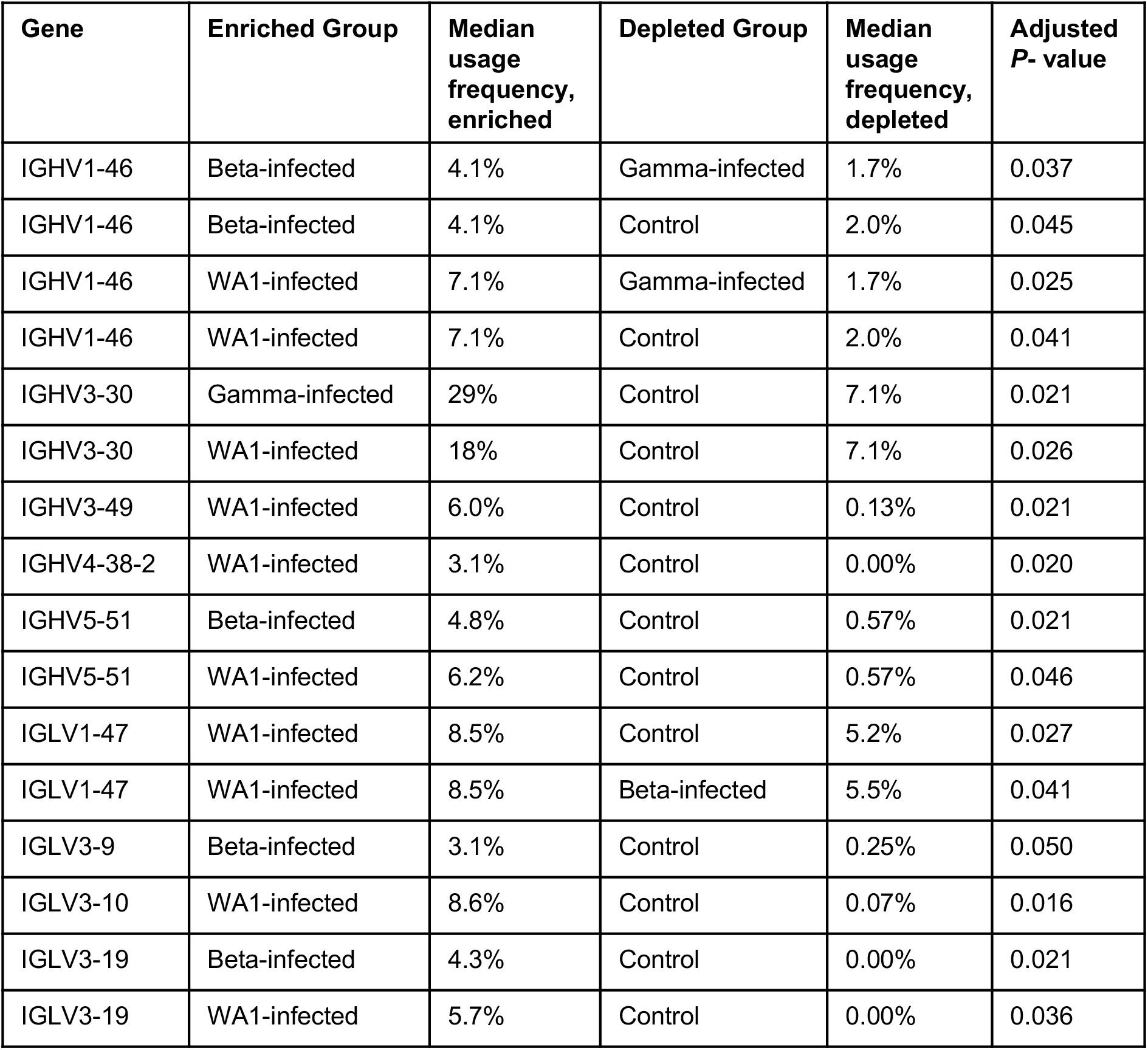
Significant differences in gene-usage. For genes with a significant difference detected by the Kruskal-Wallis test (Fig. 3C and Extended Data Fig. 7), the Dunn test was used to find significant pairwise difference. P values were adjusted for multiple testing using the Benjami-Hochberg procedure.

## Supplementary Materials

**Table 1**: Complete RATP-Ig ELISA results for SAV1. Values are reported as absorbance at 450nm wavelength (not quantitative).

**Table 2**: Complete RATP-Ig ELISA results for SAV3. Values are reported as absorbance at 450nm wavelength (not quantitative).

**Table 3**: Complete RATP-Ig ELISA results for A49. Values are reported as absorbance at 450nm wavelength (not quantitative).

**Table 4:** Sequences of peptides included in Spike pools A and B used for T cell stimulation. Highlighted peptides did not meet >75% purity and were not included in the pool.

**Table 5:** Sequences of peptides included in selected peptide pools for each variant used for T cell stimulation.

## References and Notes

1. Weisblum, Y., Schmidt, F., Zhang, F., DaSilva, J., Poston, D., Lorenzi, J.C., Muecksch, F., Rutkowska, M., Hoffmann, H.H., Michailidis, E., Gaebler, C., Agudelo, M., Cho, A., Wang, Z., Gazumyan, A., Cipolla, M., Luchsinger, L., Hillyer, C.D., Caskey, M., Robbiani, D.F., Rice, C.M., Nussenzweig, M.C., Hatziioannou, T. & Bieniasz, P.D. Escape from neutralizing antibodies by SARS-CoV-2 spike protein variants. Elife 9 (2020).

2. Schmidt, F., Weisblum, Y., Rutkowska, M., Poston, D., DaSilva, J., Zhang, F., Bednarski, E., Cho, A., Schaefer-Babajew, D.J., Gaebler, C., Caskey, M., Nussenzweig, M.C., Hatziioannou, T. & Bieniasz, P.D. High genetic barrier to SARS-CoV-2 polyclonal neutralizing antibody escape. Nature 600, 512–516 (2021).

3. Tarke, A., Coelho, C.H., Zhang, Z., Dan, J.M., Yu, E.D., Methot, N., Bloom, N.I., Goodwin, B., Phillips, E., Mallal, S., Sidney, J., Filaci, G., Weiskopf, D., da Silva Antunes, R., Crotty, S., Grifoni, A. & Sette, A. SARS-CoV-2 vaccination induces immunological T cell memory able to cross-recognize variants from Alpha to Omicron. Cell 185, 847–859 e811 (2022).

4. Pegu, A., O’Connell, S.E., Schmidt, S.D., O’Dell, S., Talana, C.A., Lai, L., Albert, J., Anderson, E., Bennett, H., Corbett, K.S., Flach, B., Jackson, L., Leav, B., Ledgerwood, J.E., Luke, C.J., Makowski, M., Nason, M.C., Roberts, P.C., Roederer, M., Rebolledo, P.A., Rostad, C.A., Rouphael, N.G., Shi, W., Wang, L., Widge, A.T., Yang, E.S. m R.N.A.S.G.s.s., Beigel, J.H., Graham, B.S., Mascola, J.R., Suthar, M.S., McDermott, A.B., Doria-Rose, N.A., Arega, J., Beigel, J.H., Buchanan, W., Elsafy, M., Hoang, B., Lampley, R., Kolhekar, A., Koo, H., Luke, C., Makhene, M., Nayak, S., Pikaart-Tautges, R., Roberts, P.C., Russell, J., Sindall, E., Albert, J., Kunwar, P., Makowski, M., Anderson, E.J., Bechnak, A., Bower, M., Camacho-Gonzalez, A.F., Collins, M., Drobeniuc, A., Edara, V.V., Edupuganti, S., Floyd, K., Gibson, T., Ackerley, C.M.G., Johnson, B., Kamidani, S., Kao, C., Kelley, C., Lai, L., Macenczak, H., McCullough, M.P., Peters, E., Phadke, V.K., Rebolledo, P.A., Rostad, C.A., Rouphael, N., Scherer, E., Sherman, A., Stephens, K., Suthar, M.S., Teherani, M., Traenkner, J., Winston, J., Yildirim, I., Barr, L., Benoit, J., Carste, B., Choe, J., Dunstan, M., Erolin, R., Ffitch, J., Fields, C., Jackson, L.A., Kiniry, E., Lasicka, S., Lee, S., Nguyen, M., Pimienta, S., Suyehira, J., Witte, M., Bennett, H., Altaras, N.E., Carfi, A., Hurley, M., Leav, B., Pajon, R., Sun, W., Zaks, T., Coler, R.N., Larsen, S.E., Neuzil, K.M., Lindesmith, L.C., Martinez, D.R., Munt, J., Mallory, M., Edwards, C., Baric, R.S., Berkowitz, N.M., Boritz, E.A., Carlton, K., Corbett, K.S., Costner, P., Creanga, A., Doria-Rose, N.A., Douek, D.C., Flach, B., Gaudinski, M., Gordon, I., Graham, B.S., Holman, L., Ledgerwood, J.E., Leung, K., Lin, B.C., Louder, M.K., Mascola, J.R., McDermott, A.B., Morabito, K.M., Novik, L., O’Connell, S., O’Dell, S., Padilla, M., Pegu, A., Schmidt, S.D., Shi, W., Swanson, P.A., 2nd, Talana, C.A., Wang, L., Widge, A.T., Yang, E.S., Zhang, Y., Chappell, J.D., Denison, M.R., Hughes, T., Lu, X., Pruijssers, A.J., Stevens, L.J., Posavad, C.M., Gale, M., Jr., Menachery, V. & Shi, P.Y. Durability of mRNA-1273 vaccine-induced antibodies against SARS-CoV-2 variants. Science 373, 1372–1377 (2021).

5. Richardson, S.I., Manamela, N.P., Motsoeneng, B.M., Kaldine, H., Ayres, F., Makhado, Z., Mennen, M., Skelem, S., Williams, N., Sullivan, N.J., Misasi, J., Gray, G.G., Bekker, L.-G., Ueckermann, V., Rossouw, T.M., Boswell, M.T., Ntusi, N.A.B., Burgers, W.A. & Moore, P.L. SARS-CoV-2 Beta and Delta variants trigger Fc effector function with increased cross-reactivity. Cell Reports Medicine 3, 100510 (2022).

6. Altarawneh, H.N., Chemaitelly, H., Ayoub, H.H., Tang, P., Hasan, M.R., Yassine, H.M., Al-Khatib, H.A., Smatti, M.K., Coyle, P., Al-Kanaani, Z., Al-Kuwari, E., Jeremijenko, A., Kaleeckal, A.H., Latif, A.N., Shaik, R.M., Abdul-Rahim, H.F., Nasrallah, G.K., Al-Kuwari, M.G., Butt, A.A., Al-Romaihi, H.E., Al-Thani, M.H., Al-Khal, A., Bertollini, R. & Abu-Raddad, L.J. Effects of Previous Infection and Vaccination on Symptomatic Omicron Infections. New England Journal of Medicine (2022).

7. Tong, P., Gautam, A., Windsor, I.W., Travers, M., Chen, Y., Garcia, N., Whiteman, N.B., McKay, L.G.A., Storm, N., Malsick, L.E., Honko, A.N., Lelis, F.J.N., Habibi, S., Jenni, S., Cai, Y., Rennick, L.J., Duprex, W.P., McCarthy, K.R., Lavine, C.L., Zuo, T., Lin, J., Zuiani, A., Feldman, J., MacDonald, E.A., Hauser, B.M., Griffths, A., Seaman, M.S., Schmidt, A.G., Chen, B., Neuberg, D., Bajic, G., Harrison, S.C. & Wesemann, D.R. Memory B cell repertoire for recognition of evolving SARS-CoV-2 spike. Cell 184, 4969–4980 e4915 (2021).

8. Greaney, A.J., Starr, T.N., Gilchuk, P., Zost, S.J., Binshtein, E., Loes, A.N., Hilton, S.K., Huddleston, J., Eguia, R., Crawford, K.H.D., Dingens, A.S., Nargi, R.S., Sutton, R.E., Suryadevara, N., Rothlauf, P.W., Liu, Z., Whelan, S.P.J., Carnahan, R.H., Crowe, J.E., Jr. & Bloom, J.D. Complete Mapping of Mutations to the SARS-CoV-2 Spike Receptor-Binding Domain that Escape Antibody Recognition. Cell Host Microbe 29, 44–57 e49 (2021).

9. Brouwer, P.J.M., Caniels, T.G., van der Straten, K., Snitselaar, J.L., Aldon, Y., Bangaru, S., Torres, J.L., Okba, N.M.A., Claireaux, M., Kerster, G., Bentlage, A.E.H., van Haaren, M.M., Guerra, D., Burger, J.A., Schermer, E.E., Verheul, K.D., van der Velde, N., van der Kooi, A., van Schooten, J., van Breemen, M.J., Bijl, T.P.L., Sliepen, K., Aartse, A., Derking, R., Bontjer, I., Kootstra, N.A., Wiersinga, W.J., Vidarsson, G., Haagmans, B.L., Ward, A.B., de Bree, G.J., Sanders, R.W. & van Gils, M.J. Potent neutralizing antibodies from COVID-19 patients define multiple targets of vulnerability. Science 369, 643–650 (2020).

10. Liu, L., Wang, P., Nair, M.S., Yu, J., Rapp, M., Wang, Q., Luo, Y., Chan, J.F., Sahi, V., Figueroa, A., Guo, X.V., Cerutti, G., Bimela, J., Gorman, J., Zhou, T., Chen, Z., Yuen, K.Y., Kwong, P.D., Sodroski, J.G., Yin, M.T., Sheng, Z., Huang, Y., Shapiro, L. & Ho, D.D. Potent neutralizing antibodies against multiple epitopes on SARS-CoV-2 spike. Nature 584, 450–456 (2020).

11. Rapp, M., Guo, Y., Reddem, E.R., Yu, J., Liu, L., Wang, P., Cerutti, G., Katsamba, P., Bimela, J.S., Bahna, F.A., Mannepalli, S.M., Zhang, B., Kwong, P.D., Huang, Y., Ho, D.D., Shapiro, L. & Sheng, Z. Modular basis for potent SARS-CoV-2 neutralization by a prevalent VH1-2-derived antibody class. Cell Rep 35, 108950 (2021).

12. Cao, Y., Su, B., Guo, X., Sun, W., Deng, Y., Bao, L., Zhu, Q., Zhang, X., Zheng, Y., Geng, C., Chai, X., He, R., Li, X., Lv, Q., Zhu, H., Deng, W., Xu, Y., Wang, Y., Qiao, L., Tan, Y., Song, L., Wang, G., Du, X., Gao, N., Liu, J., Xiao, J., Su, X.D., Du, Z., Feng, Y., Qin, C., Qin, C., Jin, R. & Xie, X.S. Potent Neutralizing Antibodies against SARS-CoV-2 Identified by High-Throughput Single-Cell Sequencing of Convalescent Patients’ B Cells. Cell 182, 73–84 e16 (2020).

13. Robbiani, D.F., Gaebler, C., Muecksch, F., Lorenzi, J.C.C., Wang, Z., Cho, A., Agudelo, M., Barnes, C.O., Gazumyan, A., Finkin, S., Hagglof, T., Oliveira, T.Y., Viant, C., Hurley, A., Hoffmann, H.H., Millard, K.G., Kost, R.G., Cipolla, M., Gordon, K., Bianchini, F., Chen, S.T., Ramos, V., Patel, R., Dizon, J., Shimeliovich, I., Mendoza, P., Hartweger, H., Nogueira, L., Pack, M., Horowitz, J., Schmidt, F., Weisblum, Y., Michailidis, E., Ashbrook, A.W., Waltari, E., Pak, J.E., Huey-Tubman, K.E., Koranda, N., Hoffman, P.R., West, A.P., Jr., Rice, C.M., Hatziioannou, T., Bjorkman, P.J., Bieniasz, P.D., Caskey, M. & Nussenzweig, M.C. Convergent antibody responses to SARS-CoV-2 in convalescent individuals. Nature 584, 437–442 (2020).

14. Cerutti, G., Guo, Y., Zhou, T., Gorman, J., Lee, M., Rapp, M., Reddem, E.R., Yu, J., Bahna, F., Bimela, J., Huang, Y., Katsamba, P.S., Liu, L., Nair, M.S., Rawi, R., Olia, A.S., Wang, P., Zhang, B., Chuang, G.Y., Ho, D.D., Sheng, Z., Kwong, P.D. & Shapiro, L. Potent SARS-CoV-2 neutralizing antibodies directed against spike N-terminal domain target a single supersite. Cell Host Microbe 29, 819–833 e817 (2021).

15. Wang, L., Zhou, T., Zhang, Y., Yang, E.S., Schramm, C.A., Shi, W., Pegu, A., Oloniniyi, O.K., Henry, A.R., Darko, S., Narpala, S.R., Hatcher, C., Martinez, D.R., Tsybovsky, Y., Phung, E., Abiona, O.M., Antia, A., Cale, E.M., Chang, L.A., Choe, M., Corbett, K.S., Davis, R.L., DiPiazza, A.T., Gordon, I.J., Hait, S.H., Hermanus, T., Kgagudi, P., Laboune, F., Leung, K., Liu, T., Mason, R.D., Nazzari, A.F., Novik, L., O’Connell, S., O’Dell, S., Olia, A.S., Schmidt, S.D., Stephens, T., Stringham, C.D., Talana, C.A., Teng, I.T., Wagner, D.A., Widge, A.T., Zhang, B., Roederer, M., Ledgerwood, J.E., Ruckwardt, T.J., Gaudinski, M.R., Moore, P.L., Doria-Rose, N.A., Baric, R.S., Graham, B.S., McDermott, A.B., Douek, D.C., Kwong, P.D., Mascola, J.R., Sullivan, N.J. & Misasi, J. Ultrapotent antibodies against diverse and highly transmissible SARS-CoV-2 variants. Science 373 (2021).

16. Moyo-Gwete, T., Madzivhandila, M., Makhado, Z., Ayres, F., Mhlanga, D., Oosthuysen, B., Lambson, B.E., Kgagudi, P., Tegally, H., Iranzadeh, A., Doolabh, D., Tyers, L., Chinhoyi, L.R., Mennen, M., Skelem, S., Marais, G., Wibmer, C.K., Bhiman, J.N., Ueckermann, V., Rossouw, T., Boswell, M., de Oliveira, T., Williamson, C., Burgers, W.A., Ntusi, N., Morris, L. & Moore, P.L. Cross-Reactive Neutralizing Antibody Responses Elicited by SARS-CoV-2 501Y.V2 (B.1.351). N Engl J Med 384, 2161–2163 (2021).

17. Liu, C., Ginn, H.M., Dejnirattisai, W., Supasa, P., Wang, B., Tuekprakhon, A., Nutalai, R., Zhou, D., Mentzer, A.J., Zhao, Y., Duyvesteyn, H.M.E., Lopez-Camacho, C., Slon-Campos, J., Walter, T.S., Skelly, D., Johnson, S.A., Ritter, T.G., Mason, C., Costa Clemens, S.A., Gomes Naveca, F., Nascimento, V., Nascimento, F., Fernandes da Costa, C., Resende, P.C., Pauvolid-Correa, A., Siqueira, M.M., Dold, C., Temperton, N., Dong, T., Pollard, A.J., Knight, J.C., Crook, D., Lambe, T., Clutterbuck, E., Bibi, S., Flaxman, A., Bittaye, M., Belij-Rammerstorfer, S., Gilbert, S.C., Malik, T., Carroll, M.W., Klenerman, P., Barnes, E., Dunachie, S.J., Baillie, V., Serafin, N., Ditse, Z., Da Silva, K., Paterson, N.G., Williams, M.A., Hall, D.R., Madhi, S., Nunes, M.C., Goulder, P., Fry, E.E., Mongkolsapaya, J., Ren, J., Stuart, D.I. & Screaton, G.R. Reduced neutralization of SARS-CoV-2 B.1.617 by vaccine and convalescent serum. Cell 184, 4220–4236 e4213 (2021).

18. Corbett, K.S., Gagne, M., Wagner, D.A., s, O.C., Narpala, S.R., Flebbe, D.R., Andrew, S.F., Davis, R.L., Flynn, B., Johnston, T.S., Stringham, C.D., Lai, L., Valentin, D., Van Ry, A., Flinchbaugh, Z., Werner, A.P., Moliva, J.I., Sriparna, M., O’Dell, S., Schmidt, S.D., Tucker, C., Choi, A., Koch, M., Bock, K.W., Minai, M., Nagata, B.M., Alvarado, G.S., Henry, A.R., Laboune, F., Schramm, C.A., Zhang, Y., Yang, E.S., Wang, L., Choe, M., Boyoglu-Barnum, S., Wei, S., Lamb, E., Nurmukhambetova, S.T., Provost, S.J., Donaldson, M.M., Marquez, J., Todd, J.M., Cook, A., Dodson, A., Pekosz, A., Boritz, E., Ploquin, A., Doria-Rose, N., Pessaint, L., Andersen, H., Foulds, K.E., Misasi, J., Wu, K., Carfi, A., Nason, M.C., Mascola, J., Moore, I.N., Edwards, D.K., Lewis, M.G., Suthar, M.S., Roederer, M., McDermott, A., Douek, D.C., Sullivan, N.J., Graham, B.S. & Seder, R.A. Protection against SARS-CoV-2 Beta variant in mRNA-1273 vaccine-boosted nonhuman primates. Science 374, 1343–1353 (2021).

19. Gagne, M.M. J.I.; Foulds, K.E.; Andrew, S.F.; Flynn, B.J.; Werner, A.P.; Wagner, D.A.; Teng, I.-T.; Lin, B.C.; Moore, C.; Jean-Baptiste, N.; Carroll, R.; Foster, S.L.; Patel, M.; Ellis, M.; Edara, V.-V.; Maldonado, N.V.; Minai, M.; McCormick, L.; Honeycutt, C.C.; Nagata, B.M.; Bock, K.W.; Dulan, C.N.M.; Cordon, J.; Flebbe, D.R.; Todd, J.-P.M.; McCarthy, E.; Pessaint, L.; Van Ry, A.; Narvaez, B.; Valentin, D.; Cook, A.; Dodson, A.; Steingrebe, K.; Nurmukhambetova, S.T.; & Godbole, S.H. A.R.; Laboune, F.; Roberts-Torres, J.; Lorang, C.G.; Amin, S.; Trost, J.; Naisan, M.; Basappa, M.; Willis, J.; Wang, L.; Shi, W.; Doria-Rose, N.A.; Zhang, Y.; Yang, E.S.; Leung, K.; O’Dell, S.; Schmidt, S.D.; Olia, A.S.; Liu, C.; Harris, D.R.; Chuang, G.-Y.; Stewart-Jones, G.; Renzi, I.; Lai, Y.-T.; Malinowski, A.; Wu, K.; Mascola, J.R.; Carfi, A.; Kwong, P.D.; Edwards, D.K.; Lewis, M.G.; Andersen, H.; Corbett, K.S.; Nason, M.C.; McDermott, A.B.; Suthar, M.S.; Moore, I.N.; Roederer, M.; Sullivan, N.J.; Douek, D.C.; Seder, R.A. mRNA-1273 or mRNA-Omicron boost in vaccinated macaques elicits comparable B cell expansion, neutralizing antibodies and protection against Omicron. Cell (2022).

20. Li, D., Edwards, R.J., Manne, K., Martinez, D.R., Schafer, A., Alam, S.M., Wiehe, K., Lu, X., Parks, R., Sutherland, L.L., Oguin, T.H., 3rd, McDanal, C., Perez, L.G., Mansouri, K., Gobeil, S.M.C., Janowska, K., Stalls, V., Kopp, M., Cai, F., Lee, E., Foulger, A., Hernandez, G.E., Sanzone, A., Tilahun, K., Jiang, C., Tse, L.V., Bock, K.W., Minai, M., Nagata, B.M., Cronin, K., Gee-Lai, V., Deyton, M., Barr, M., Von Holle, T., Macintyre, A.N., Stover, E., Feldman, J., Hauser, B.M., Caradonna, T.M., Scobey, T.D., Rountree, W., Wang, Y., Moody, M.A., Cain, D.W., DeMarco, C.T., Denny, T.N., Woods, C.W., Petzold, E.W., Schmidt, A.G., Teng, I.T., Zhou, T., Kwong, P.D., Mascola, J.R., Graham, B.S., Moore, I.N., Seder, R., Andersen, H., Lewis, M.G., Montefiori, D.C., Sempowski, G.D., Baric, R.S., Acharya, P., Haynes, B.F. & Saunders, K.O. In vitro and in vivo functions of SARS-CoV-2 infection-enhancing and neutralizing antibodies. Cell 184, 4203–4219 e4232 (2021).

21. Wec, A.Z., Wrapp, D., Herbert, A.S., Maurer, D.P., Haslwanter, D., Sakharkar, M., Jangra, R.K., Dieterle, M.E., Lilov, A., Huang, D., Tse, L.V., Johnson, N.V., Hsieh, C.L., Wang, N., Nett, J.H., Champney, E., Burnina, I., Brown, M., Lin, S., Sinclair, M., Johnson, C., Pudi, S., Bortz, R., 3rd, Wirchnianski, A.S., Laudermilch, E., Florez, C., Fels, J.M., O’Brien, C.M., Graham, B.S., Nemazee, D., Burton, D.R., Baric, R.S., Voss, J.E., Chandran, K., Dye, J.M., McLellan, J.S. & Walker, L.M. Broad neutralization of SARS-related viruses by human monoclonal antibodies. Science 369, 731–736 (2020).

22. Zost, S.J., Gilchuk, P., Chen, R.E., Case, J.B., Reidy, J.X., Trivette, A., Nargi, R.S., Sutton, R.E., Suryadevara, N., Chen, E.C., Binshtein, E., Shrihari, S., Ostrowski, M., Chu, H.Y., Didier, J.E., MacRenaris, K.W., Jones, T., Day, S., Myers, L., Eun-Hyung Lee, F., Nguyen, D.C., Sanz, I., Martinez, D.R., Rothlauf, P.W., Bloyet, L.M., Whelan, S.P.J., Baric, R.S., Thackray, L.B., Diamond, M.S., Carnahan, R.H. & Crowe, J.E., Jr. Rapid isolation and profiling of a diverse panel of human monoclonal antibodies targeting the SARS-CoV-2 spike protein. Nat Med 26, 1422–1427 (2020).

23. Song, G., He, W.T., Callaghan, S., Anzanello, F., Huang, D., Ricketts, J., Torres, J.L., Beutler, N., Peng, L., Vargas, S., Cassell, J., Parren, M., Yang, L., Ignacio, C., Smith, D.M., Voss, J.E., Nemazee, D., Ward, A.B., Rogers, T., Burton, D.R. & Andrabi, R. Cross-reactive serum and memory B-cell responses to spike protein in SARS-CoV-2 and endemic coronavirus infection. Nat Commun 12, 2938 (2021).

24. Liu, L., Iketani, S., Guo, Y., Reddem, E.R., Casner, R.G., Nair, M.S., Yu, J., Chan, J.F.-W., Wang, M., Cerutti, G., Li, Z., Morano, N.C., Castagna, C.D., Corredor, L., Chu, H., Yuan, S., Poon, V.K.-M., Chan, C.C.-S., Chen, Z., Luo, Y., Cunningham, M., Chavez, A., Yin, M.T., Perlin, D.S., Tsuji, M., Yuen, K.-Y., Kwong, P.D., Sheng, Z., Huang, Y., Shapiro, L. & Ho, D.D. An antibody class with a common CDRH3 motif broadly neutralizes sarbecoviruses. Science Translational Medicine 14, eabn6859 (2022).

25. Soto, C., Bombardi, R.G., Branchizio, A., Kose, N., Matta, P., Sevy, A.M., Sinkovits, R.S., Gilchuk, P., Finn, J.A. & Crowe, J.E., Jr. High frequency of shared clonotypes in human B cell receptor repertoires. Nature 566, 398–402 (2019).

26. Liu, C., Zhou, D., Nutalai, R., Duyvesteyn, H.M.E., Tuekprakhon, A., Ginn, H.M., Dejnirattisai, W., Supasa, P., Mentzer, A.J., Wang, B., Case, J.B., Zhao, Y., Skelly, D.T., Chen, R.E., Johnson, S.A., Ritter, T.G., Mason, C., Malik, T., Temperton, N., Paterson, N.G., Williams, M.A., Hall, D.R., Clare, D.K., Howe, A., Goulder, P.J.R., Fry, E.E., Diamond, M.S., Mongkolsapaya, J., Ren, J., Stuart, D.I. & Screaton, G.R. The antibody response to SARS-CoV-2 Beta underscores the antigenic distance to other variants. Cell Host Microbe 30, 53–68 e12 (2022).

27. Reincke, S.M., Yuan, M., Kornau, H.C., Corman, V.M., van Hoof, S., Sanchez-Sendin, E., Ramberger, M., Yu, W., Hua, Y., Tien, H., Schmidt, M.L., Schwarz, T., Jeworowski, L.M., Brandl, S.E., Rasmussen, H.F., Homeyer, M.A., Stoffler, L., Barner, M., Kunkel, D., Huo, S., Horler, J., von Wardenburg, N., Kroidl, I., Eser, T.M., Wieser, A., Geldmacher, C., Hoelscher, M., Ganzer, H., Weiss, G., Schmitz, D., Drosten, C., Pruss, H., Wilson, I.A. & Kreye, J. SARS-CoV-2 Beta variant infection elicits potent lineage-specific and cross-reactive antibodies. Science 375, 782–787 (2022).

28. Seydoux, E., Homad, L.J., MacCamy, A.J., Parks, K.R., Hurlburt, N.K., Jennewein, M.F., Akins, N.R., Stuart, A.B., Wan, Y.H., Feng, J., Whaley, R.E., Singh, S., Boeckh, M., Cohen, K.W., McElrath, M.J., Englund, J.A., Chu, H.Y., Pancera, M., McGuire, A.T. & Stamatatos, L. Analysis of a SARS-CoV-2-Infected Individual Reveals Development of Potent Neutralizing Antibodies with Limited Somatic Mutation. Immunity 53, 98–105 e105 (2020).

29. Wang, Y., Yuan, M., Lv, H., Peng, J., Wilson, I.A. & Wu, N.C. A large-scale systematic survey reveals recurring molecular features of public antibody responses to SARS-CoV-2. Immunity 55, 1105-1117.e1104 (2022).

30. Dussupt, V., Sankhala, R.S., Mendez-Rivera, L., Townsley, S.M., Schmidt, F., Wieczorek, L., Lal, K.G., Donofrio, G.C., Tran, U., Jackson, N.D., Zaky, W.I., Zemil, M., Tritsch, S.R., Chen, W.H., Martinez, E.J., Ahmed, A., Choe, M., Chang, W.C., Hajduczki, A., Jian, N., Peterson, C.E., Rees, P.A., Rutkowska, M., Slike, B.M., Selverian, C.N., Swafford, I., Teng, I.T., Thomas, P.V., Zhou, T., Smith, C.J., Currier, J.R., Kwong, P.D., Rolland, M., Davidson, E., Doranz, B.J., Mores, C.N., Hatziioannou, T., Reiley, W.W., Bieniasz, P.D., Paquin-Proulx, D., Gromowski, G.D., Polonis, V.R., Michael, N.L., Modjarrad, K., Joyce, M.G. & Krebs, S.J. Low-dose in vivo protection and neutralization across SARS-CoV-2 variants by monoclonal antibody combinations. Nat Immunol 22, 1503–1514 (2021).

31. Wang, P., Nair, M.S., Liu, L., Iketani, S., Luo, Y., Guo, Y., Wang, M., Yu, J., Zhang, B., Kwong, P.D., Graham, B.S., Mascola, J.R., Chang, J.Y., Yin, M.T., Sobieszczyk, M., Kyratsous, C.A., Shapiro, L., Sheng, Z., Huang, Y. & Ho, D.D. Antibody resistance of SARS-CoV-2 variants B.1.351 and B.1.1.7. Nature 593, 130–135 (2021).

32. Laurie, M.T., Liu, J., Sunshine, S., Peng, J., Black, D., Mitchell, A.M., Mann, S.A., Pilarowski, G., Zorn, K.C., Rubio, L., Bravo, S., Marquez, C., Sabatino, J.J., Jr, Mittl, K., Petersen, M., Havlir, D. & DeRisi, J. SARS-CoV-2 Variant Exposures Elicit Antibody Responses With Differential Cross-Neutralization of Established and Emerging Strains Including Delta and Omicron. The Journal of Infectious Diseases 225, 1909–1914 (2022).

33. Greaney, A.J., Starr, T.N., Eguia, R.T., Loes, A.N., Khan, K., Karim, F., Cele, S., Bowen, J.E., Logue, J.K., Corti, D., Veesler, D., Chu, H.Y., Sigal, A. & Bloom, J.D. A SARS-CoV-2 variant elicits an antibody response with a shifted immunodominance hierarchy. PLoS Pathog 18, e1010248 (2022).

34. Nutalai, R., Zhou, D., Tuekprakhon, A., Ginn, H.M., Supasa, P., Liu, C., Huo, J., Mentzer, A.J., Duyvesteyn, H.M.E., Dijokaite-Guraliuc, A., Skelly, D., Ritter, T.G., Amini, A., Bibi, S., Adele, S., Johnson, S.A., Constantinides, B., Webster, H., Temperton, N., Klenerman, P., Barnes, E., Dunachie, S.J., Crook, D., Pollard, A.J., Lambe, T., Goulder, P., Paterson, N.G., Williams, M.A., Hall, D.R., Mongkolsapaya, J., Fry, E.E., Dejnirattisai, W., Ren, J., Stuart, D.I. & Screaton, G.R. Potent cross-reactive antibodies following Omicron breakthrough in vaccinees. Cell 185, 2116-2131.e2118 (2022).

35. Cao, Y., Yisimayi, A., Jian, F., Song, W., Xiao, T., Wang, L., Du, S., Wang, J., Li, Q., Chen, X., Wang, P., Zhang, Z., Liu, P., An, R., Hao, X., Wang, Y., Wang, J., Feng, R., Sun, H., Zhao, L., Zhang, W., Zhao, D., Zheng, J., Yu, L., Li, C., Zhang, N., Wang, R., Niu, X., Yang, S., Song, X., Zheng, L., Li, Z., Gu, Q., Shao, F., Huang, W., Jin, R., Shen, Z., Wang, Y., Wang, X., Xiao, J. & Xie, X.S. BA.2.12.1, BA.4 and BA.5 escape antibodies elicited by Omicron infection. bioRxiv, 2022.2004.2030.489997 (2022).

36. Reynolds, C.J., Pade, C., Gibbons, J.M., Otter, A.D., Lin, K.-M., Sandoval, D.M., Pieper, F.P., Butler, D.K., Liu, S., Joy, G., Forooghi, N., Treibel, T.A., Manisty, C., Moon, J.C., Semper, A., Brooks, T., McKnight, Á., Altmann, D.M., Boyton, R.J., Abbass, H., Abiodun, A., Alfarih, M., Alldis, Z., Altmann, D.M., Amin, O.E., Andiapen, M., Artico, J., Augusto, J.B., Baca, G.L., Bailey, S.N.L., Bhuva, A.N., Boulter, A., Bowles, R., Boyton, R.J., Bracken, O.V., O’Brien, B., Brooks, T., Bullock, N., Butler, D.K., Captur, G., Carr, O., Champion, N., Chan, C., Chandran, A., Coleman, T., Sousa, J.C.d., Couto-Parada, X., Cross, E., Cutino-Moguel, T., D’Arcangelo, S., Davies, R.H., Douglas, B., Genova, C.D., Dieobi-Anene, K., Diniz, M.O., Ellis, A., Feehan, K., Finlay, M., Fontana, M., Forooghi, N., Francis, S., Gibbons, J.M., Gillespie, D., Gilroy, D., Hamblin, M., Harker, G., Hemingway, G., Hewson, J., Heywood, W., Hickling, L.M., Hicks, B., Hingorani, A.D., Howes, L., Itua, I., Jardim, V., Lee, W.-Y.J., Jensen, M., Jones, J., Jones, M., Joy, G., Kapil, V., Kelly, C., Kurdi, H., Lambourne, J., Lin, K.-M., Liu, S., Lloyd, A., Louth, S., Maini, M.K., Mandadapu, V., Manisty, C., McKnight, Á., Menacho, K., Mfuko, C., Mills, K., Millward, S., Mitchelmore, O., Moon, C., Moon, J., Sandoval, D.M., Murray, S.M., Noursadeghi, M., Otter, A., Pade, C., Palma, S., Parker, R., Patel, K., Pawarova, M., Petersen, S.E., Piniera, B., Pieper, F.P., Rannigan, L., Rapala, A., Reynolds, C.J., Richards, A., Robathan, M., Rosenheim, J., Rowe, C., Royds, M., West, J.S., Sambile, G., Schmidt, N.M., Selman, H., Semper, A., Seraphim, A., Simion, M., Smit, A., Sugimoto, M., Swadling, L., Taylor, S., Temperton, N., Thomas, S., Thornton, G.D., Treibel, T.A., Tucker, A., Varghese, A., Veerapen, J., Vijayakumar, M., Warner, T., Welch, S., White, H., Wodehouse, T., Wynne, L., Zahedi, D., Chain, B. & Moon, J.C. Immune boosting by B.1.1.529 <b>(</b>Omicron) depends on previous SARS-CoV-2 exposure. Science 0, eabq1841.

37. Beaudoin-Bussieres, G., Chen, Y., Ullah, I., Prevost, J., Tolbert, W.D., Symmes, K., Ding, S., Benlarbi, M., Gong, S.Y., Tauzin, A., Gasser, R., Chatterjee, D., Vezina, D., Goyette, G., Richard, J., Zhou, F., Stamatatos, L., McGuire, A.T., Charest, H., Roger, M., Pozharski, E., Kumar, P., Mothes, W., Uchil, P.D., Pazgier, M. & Finzi, A. A Fc-enhanced NTD-binding non-neutralizing antibody delays virus spread and synergizes with a nAb to protect mice from lethal SARS-CoV-2 infection. Cell Rep 38, 110368 (2022).

38. Robinson, S.A., Raybould, M.I.J., Schneider, C., Wong, W.K., Marks, C. & Deane, C.M. Epitope profiling using computational structural modelling demonstrated on coronavirus-binding antibodies. PLoS Comput Biol 17, e1009675 (2021).

39. Gagne, M., Moliva, J.I., Foulds, K.E., Andrew, S.F., Flynn, B.J., Werner, A.P., Wagner, D.A., Teng, I.T., Lin, B.C., Moore, C., Jean-Baptiste, N., Carroll, R., Foster, S.L., Patel, M., Ellis, M., Edara, V.-V., Maldonado, N.V., Minai, M., McCormick, L., Honeycutt, C.C., Nagata, B.M., Bock, K.W., Dulan, C.N.M., Cordon, J., Flebbe, D.R., Todd, J.-P.M., McCarthy, E., Pessaint, L., Van Ry, A., Narvaez, B., Valentin, D., Cook, A., Dodson, A., Steingrebe, K., Nurmukhambetova, S.T., Godbole, S., Henry, A.R., Laboune, F., Roberts-Torres, J., Lorang, C.G., Amin, S., Trost, J., Naisan, M., Basappa, M., Willis, J., Wang, L., Shi, W., Doria-Rose, N.A., Zhang, Y., Yang, E.S., Leung, K., O’Dell, S., Schmidt, S.D., Olia, A.S., Liu, C., Harris, D.R., Chuang, G.-Y., Stewart-Jones, G., Renzi, I., Lai, Y.-T., Malinowski, A., Wu, K., Mascola, J.R., Carfi, A., Kwong, P.D., Edwards, D.K., Lewis, M.G., Andersen, H., Corbett, K.S., Nason, M.C., McDermott, A.B., Suthar, M.S., Moore, I.N., Roederer, M., Sullivan, N.J., Douek, D.C. & Seder, R.A. mRNA-1273 or mRNA-Omicron boost in vaccinated macaques elicits similar B cell expansion, neutralizing responses, and protection from Omicron. Cell 185, 1556-1571.e1518 (2022).

40. Ying, B., Scheaffer, S.M., Whitener, B., Liang, C.-Y., Dmytrenko, O., Mackin, S., Wu, K., Lee, D., Avena, L.E., Chong, Z., Case, J.B., Ma, L., Kim, T.T.M., Sein, C.E., Woods, A., Berrueta, D.M., Chang, G.-Y., Stewart-Jones, G., Renzi, I., Lai, Y.-T., Malinowski, A., Carfi, A., Elbashir, S.M., Edwards, D.K., Thackray, L.B. & Diamond, M.S. Boosting with variant-matched or historical mRNA vaccines protects against Omicron infection in mice. Cell 185, 1572-1587.e1511 (2022).

41. Wilks, S.H., Mühlemann, B., Shen, X., Türeli, S., LeGresley, E.B., Netzl, A., Caniza, M.A., Chacaltana-Huarcaya, J.N., Daniell, X., Datto, M.B., Denny, T.N., Drosten, C., Fouchier, R.A.M., Garcia, P.J., Halfmann, P.J., Jassem, A., Jones, T.C., Kawaoka, Y., Krammer, F., McDanal, C., Pajon, R., Simon, V., Stockwell, M., Tang, H., van Bakel, H., Webby, R., Montefiori, D.C. & Smith, D.J. Mapping SARS-CoV-2 antigenic relationships and serological responses. bioRxiv, 2022.2001.2028.477987 (2022).

42. Naldini, L., Blomer, U., Gage, F.H., Trono, D. & Verma, I.M. Efficient transfer, integration, and sustained long-term expression of the transgene in adult rat brains injected with a lentiviral vector. Proc Natl Acad Sci U S A 93, 11382–11388 (1996).

43. Zhou, T., Teng, I.T., Olia, A.S., Cerutti, G., Gorman, J., Nazzari, A., Shi, W., Tsybovsky, Y., Wang, L., Wang, S., Zhang, B., Zhang, Y., Katsamba, P.S., Petrova, Y., Banach, B.B., Fahad, A.S., Liu, L., Lopez Acevedo, S.N., Madan, B., Oliveira de Souza, M., Pan, X., Wang, P., Wolfe, J.R., Yin, M., Ho, D.D., Phung, E., DiPiazza, A., Chang, L.A., Abiona, O.M., Corbett, K.S., DeKosky, B.J., Graham, B.S., Mascola, J.R., Misasi, J., Ruckwardt, T., Sullivan, N.J., Shapiro, L. & Kwong, P.D. Structure-Based Design with Tag-Based Purification and In-Process Biotinylation Enable Streamlined Development of SARS-CoV-2 Spike Molecular Probes. Cell Rep 33, 108322 (2020).

44. Teng, I.T., Nazzari, A.F., Choe, M., Liu, T., de Souza, M.O., Petrova, Y., Tsybovsky, Y., Wang, S., Zhang, B., Artamonov, M., Madan, B., Huang, A., Lopez Acevedo, S.N., Pan, X., Ruckwardt, T.J., DeKosky, B.J., Mascola, J.R., Misasi, J., Sullivan, N.J., Zhou, T. & Kwong, P.D. Molecular probes of spike ectodomain and its subdomains for SARS-CoV-2 variants, Alpha through Omicron. bioRxiv (2021).

45. Krebs, S.J., Kwon, Y.D., Schramm, C.A., Law, W.H., Donofrio, G., Zhou, K.H., Gift, S., Dussupt, V., Georgiev, I.S., Schatzle, S., McDaniel, J.R., Lai, Y.T., Sastry, M., Zhang, B., Jarosinski, M.C., Ransier, A., Chenine, A.L., Asokan, M., Bailer, R.T., Bose, M., Cagigi, A., Cale, E.M., Chuang, G.Y., Darko, S., Driscoll, J.I., Druz, A., Gorman, J., Laboune, F., Louder, M.K., McKee, K., Mendez, L., Moody, M.A., O’Sullivan, A.M., Owen, C., Peng, D., Rawi, R., Sanders-Buell, E., Shen, C.H., Shiakolas, A.R., Stephens, T., Tsybovsky, Y., Tucker, C., Verardi, R., Wang, K., Zhou, J., Zhou, T., Georgiou, G., Alam, S.M., Haynes, B.F., Rolland, M., Matyas, G.R., Polonis, V.R., McDermott, A.B., Douek, D.C., Shapiro, L., Tovanabutra, S., Michael, N.L., Mascola, J.R., Robb, M.L., Kwong, P.D. & Doria-Rose, N.A. Longitudinal Analysis Reveals Early Development of Three MPER-Directed Neutralizing Antibody Lineages from an HIV-1-Infected Individual. Immunity 50, 677–691 e613 (2019).

46. Upadhyay, A.A., Kauffman, R.C., Wolabaugh, A.N., Cho, A., Patel, N.B., Reiss, S.M., Havenar-Daughton, C., Dawoud, R.A., Tharp, G.K., Sanz, I., Pulendran, B., Crotty, S., Lee, F.E., Wrammert, J. & Bosinger, S.E. BALDR: a computational pipeline for paired heavy and light chain immunoglobulin reconstruction in single-cell RNA-seq data. Genome Med 10, 20 (2018).

47. Schramm, C.A., Sheng, Z., Zhang, Z., Mascola, J.R., Kwong, P.D. & Shapiro, L. SONAR: A High-Throughput Pipeline for Inferring Antibody Ontogenies from Longitudinal Sequencing of B Cell Transcripts. Front Immunol 7, 372 (2016).

48. Hao, Y., Hao, S., Andersen-Nissen, E., Mauck, W.M., 3rd, Zheng, S., Butler, A., Lee, M.J., Wilk, A.J., Darby, C., Zager, M., Hoffman, P., Stoeckius, M., Papalexi, E., Mimitou, E.P., Jain, J., Srivastava, A., Stuart, T., Fleming, L.M., Yeung, B., Rogers, A.J., McElrath, J.M., Blish, C.A., Gottardo, R., Smibert, P. & Satija, R. Integrated analysis of multimodal single-cell data. Cell 184, 3573–3587 e3529 (2021).

49. Rognes, T., Flouri, T., Nichols, B., Quince, C. & Mahe, F. VSEARCH: a versatile open source tool for metagenomics. PeerJ 4, e2584 (2016).

